# Calcium/Calmodulin Dependent Protein Kinase Kinase 2 Regulates the Expansion of Tumor-induced Myeloid-Derived Suppressor Cells

**DOI:** 10.1101/2021.07.28.454009

**Authors:** Wei Huang, Yaping Liu, Anthony Luz, Mark Berrong, Joel N Meyer, Yujing Zou, Excel Swann, Pasupathi Sundaramoorthy, Yubin Kang, Shekeab Jauhari, William Lento, Nelson Chao, Luigi Racioppi

**Affiliations:** Division of Hematological Malignancies and Cellular Therapy, Department of Medicine, Duke University Medical Center, Durham, NC 27710. USA; Duke University Nicholas School of the Environment. Durham, NC 27710. USA; Duke Human Vaccine Institute, Durham, NC 27710, USA; Department of Molecular Medicine and Medical Biotechnology, University of Naples Federico II, 80131 Naples, Italy

## Abstract

Myeloid-derived suppressor cells (MDSCs) are a heterogeneous group of cells, which can suppress the immune response, promote tumor progression and impair the efficacy of immunotherapies. Consequently, the pharmacological targeting of MDSC is emerging as a new immunotherapeutic strategy to stimulate the natural anti-tumor immune response and potentiate the efficacy of immunotherapies. Herein, we leveraged genetically modified models and a small molecule inhibitor to validate Calcium-Calmodulin Kinase Kinase 2 (CaMKK2) as a druggable target to control MDSC accumulation in tumor-bearing mice. The results indicated that deletion of CaMKK2 in the host attenuated the growth of engrafted tumor cells, and this phenomenon was associated with increased antitumor T cell response and decreased accumulation of MDSC. The adoptive transfer of MDSC was sufficient to restore the ability of the tumor to grow in *Camkk2^-/-^* mice, confirming the key role of MDSC in the mechanism of tumor rejection. In vitro studies indicated that blocking of CaMKK2 is sufficient to impair the yield of MDSC. Surprisingly, MDSC generated from *Camkk2^-/-^* bone marrow cells also showed a higher ability to terminally differentiate toward more immunogenic cell types (e.g inflammatory macrophages and dendritic cells) compared to wild type (WT). Higher intracellular levels of reactive oxygen species (ROS) accumulated in *Camkk2^-/-^* MDSC, increasing their susceptibility to apoptosis and promoting their terminal differentiation toward more mature myeloid cells. Mechanistic studies indicated that AMP-activated protein kinase (AMPK), which is a known CaMKK2 proximal target controlling the oxidative stress response, fine-tunes ROS accumulation in MDSC. Accordingly, failure to activate the CaMKK2-AMPK axis can account for the elevated ROS levels in *Camkk2^-/-^* MDSC. These results highlight CaMKK2 as an important regulator of the MDSC lifecycle, identifying this kinase as a new druggable target to restrain MDSC expansion and enhance the efficacy of anti-tumor immunotherapy.

## Introduction

Immunosuppressive myeloid cells showing the phenotype of neutrophils and monocytes were originally described more than 20 years ago and later, these cells were named myeloid-derived suppressor cells (MDSCs)^1^. For many years the features that distinguish MDSCs from conventional neutrophils and monocytes have been poorly understood.

However, leveraging genomic, proteomic, and metabolic approaches, emerging studies have contributed to the identificaton of features that distinguish MDSCs from conventional neutrophils and monocytes^2, 3^. In mice, MDSCs are phenotypically identified as CD11b^+^Gr1^+^ cells and are further categorized according to Gr-1 subgroup markers as either monocytic (CD11b+Ly6C^high^Ly6G^neg^, M-MDSC) or granulocytic MDSC (CD11b+Ly6C^low^Ly6G^high^, G-MDSC)^2, 4, 5^. In humans, M- and G-MDSCs are defined as CD11b+CD14+CD15- and CD11b+CD14-CD15+, respectively. MDSCs suppress the immune response through multiple mechanisms including the release of reactive oxygen species (ROS), nitric oxide (NO), indoleamine 2,3-dioxygenase, and suppressive cytokines^2, 6–9^. MDSCs also have the ability to deplete tissues of arginine required for T-cell proliferation. The development of MDSCs seems to involve two phases that ultimately lead to the accumulation of pathologically activated MDSCs into the tumor microenvironment (TME)^10^. During the initial phases, the tumor-derived signals drive the expansion and conditioning of bone marrow and spleen resident myeloid progenitors toward MDSCs. Subsequently, MDSCs are recruited and pathologically activated in the TME, where M-MDSC terminally differentiate into tumor-associated macrophages^2^. Overall, MDSCs function to facilitate the escape of tumor cells from the immune surveillance and the accumulation of these cell types in TME correlates with tumor stage, progression, and resistance to standard chemotherapy, radiotherapy, and immunotherapy^11–14^. For these reasons, the druggable components of the molecular machinery that controls MDSC lifecycle are becoming attractive immunotherapeutic targets to re-shape the immunosuppressive TME and enhance the efficacy of immunotherapies^15^.

One such potential pathway is calcium signaling. Calcium ions (Ca^2+^) are an important and pervasive second messenger that regulates a plethora of cell functions^16^. Most of the effects of Ca^2+^ are mediated by Calmodulin (CaM), which is a ubiquitously expressed small protein that functions as the primary intercellular Ca^2+^sensor. Once formed, the Ca^2+^/CaM complexes can bind and regulate the activity of Calcium/Calmodulin Kinase proteins (CaMKs), which is a family of Ser/Thr kinases including Ca^2+/^CaM-dependent protein kinase I (CaMKI), CaMKIV, Ca^2+/^CaM-dependent protein kinase kinase 1 (CaMKK1) and CaMKK2^17^. Following activation by Ca^2+^/CaM complexes, CaMKK1 and 2 can further phosphorylate and stimulate CaMKI and CaMKIV kinase activity. Importantly, only CAMKK2 can phosphorylate and activate 5’ AMP-activated kinase (AMPK)^18, 19^, which has an important role in the control of cellular energy metabolism, oxidative stress response, and processes that regulate innate and adaptive immunity^20, 21^. Outside the brain, CaMKK2 is expressed at detectable levels restricted to a small subset of cell types, including hematopoietic stem and progenitor cells (HSCP) and mature myeloid cell subsets such as circulating monocytes and macrophages^22^. CaMKK2 also regulates granulopoiesis and recently we have demonstrated that this kinase controls HSCP regeneration following bone marrow injury^23, 24^. Interestingly, CaMKK2 impinges the activation program of macrophages, and consequently, genetic ablation of *Camkk2* attenuates the detrimental inflammatory response induced by bacterial endotoxin or obesity^25^. More recently, we demonstrated that CaMKK2 is a key regulator of the immune-suppressive microenvironment in breast cancer and blockade of this enzyme attenuates tumor growth in a CD8+ T cell-dependent manner^26^. Hence, CaMKK2 is an important druggable target of therapies aimed to counteract tumor-induced immunosuppression and foster the anti-tumor immune response.

In this study, we investigated the role of CaMKK2 in MDSCs. To this end, we leveraged germline and inducible models of *Camkk2-*deficient mice in combination with *in vivo* and *in vitro* models of MDSC expansion. The results from these studies indicated that deletion of CaMKK2 in the host attenuated the growth of engrafted tumor cells, and this phenomenon was associated with increased antitumor T cells response and decreased accumulation of MDSCs. Mechanistically we show that the CaMKK2-AMPK axis controls the survival and the terminal differentiation of tumor-induced MDSCs by fine-tuning the accumulation of reactive oxygen species (ROS). Translationally, these results identify CaMKK2 as a novel druggable target to restrain MDSC expansion and enhance the efficacy of anti-tumor immunotherapy.

## Materials and Methods

### Mice

C57BL/6 mice were purchased from the Jackson Laboratory (Jackson Laboratory, CA, USA). CaMKK2 global knock-out (*Camkk2^-/-^*), Tg (Camkk2-EGFP)DF129Gsat mice were generated as described^27, 28^ and backcrossed to C57BL/6 for at least 8 generations. LysMCre^+^; CaMKK2^fl/fl^ and littermates control were generated by crossing B6.129P2-Lyz2tm1(cre)Ifo/J mice from the Jackson Laboratory with CaMKK2^lox/lox^ mice^29^. ERT2-Cre; CaMKK2^fl/fl^ were generated by crossing B6.Cg-Tg(CAG-cre/Esr1*)5Amc/J transgenic mice from the Jackson laboratory with CaMKK2^lox/lox^ mice^29^. All the mice were used between 8-16 weeks of age and were gender and age-matched. Animal care and experimental procedures were approved by the National Institute of Health and the Duke University Institutional Animal Care and Use Committee.

### E.G7-OVA Tumor Model

E.G7-OVA cells were purchased from American Type Tissue Culture Collection (Rockville, MA, USA). For bioluminescent analysis, E.G7-OVA cells were stably transfected with a plasmid expressing luciferase. The cells were cultured in RPMI 1640 supplied with 4.5g/L glucose, 2mM L-glutamine, 1.5g/L sodium bicarbonate, 10mM HEPES, 1.0mM sodium pyruvate (all from Gibco, MA, USA), 0.05mM 2-mercaptoethanol, 0.4mg/ml G418 (Sigma-Aldrich, MO, USA), and 10% fetal bovine serum (Hyclone, MA, USA) in humidified 37°C CO2 incubator. E.G7-OVA cells were subcutaneously injected into the flank and tumors were monitored using weekly bioluminescence imaging. The tumors were measured with a calibrator every 3-4 days and the volume was calculated as Length × Width × Width/2. EL4 T lymphoma cells (ATCC TIB-39) were purchased from Duke Cell Culture Facility and were cultured in RPMI 1640 supplied with 4.5g/L glucose, 2mM L-glutamine, 1.5g/L sodium bicarbonate, 10mM HEPES, 1.0mM sodium pyruvate (all from Gibco, MA, USA), 0.05mM 2- mercaptoethanol, and 10% fetal bovine serum (Hyclone, MA, USA) in humidified 37°C CO2 incubator. EL4 tumors were measured with a calibrator every 3-4 days and the volume was calculated as described previously. EO771^30^, 4T1^31^, MET1^32^, and A7C11^33^ epithelial mammary tumor cell lines were cultured in RPMI 1640 supplied with 4.5g/L glucose, 2mM L-glutamine, 1.5g/L sodium bicarbonate, 10mM HEPES, 1.0mM sodium pyruvate (all from Gibco, MA, USA) and 8% of 10% fetal bovine serum (Hyclone, MA, USA) in humidified 37°C CO2 incubator. To generate tumor conditioned medium (TCM), regular media from 70-80% confluent cell cultures were removed and replaced with fresh culture medium. After 72 hours TCMs were then harvested, aliquoted, and stored at −80°c before use.

### MDSC transfusion

MDSCs were isolated from the spleens of WT and *Camkk2^-/-^* mice 21 days after E.G7-OVA injection and were enriched by anti-mouse Gr-1 Microbeads (Miltenyi, Bergisch Gladbach, Germany) according to the manufacturer’s instructions. Then, 2×10^6^ MDSCs (> 90% pure) were injected into *Camkk2^-/-^* mice through the tail vein, followed by 1×10^5^ E.G7-OVA tumor cells injected subcutaneously into the flank. Tumor size was measured as described above.

### T-Cell Depletion Treatment

*Camkk2^-/-^* mice were injected peritoneally with anti-mouse CD8 monoclonal antibody or its IgG isotype (BioXCell, NH, USA) 0.2mg 4 days and 1 day before tumor inoculation, and every 3 days after EG7 tumor cell injection. CD8^+^ T cells depletion was monitored by flow cytometry.

### Flow Cytometry Analysis

Single-cell suspensions of splenocytes were prepared as described^34^. The removed tumors were dissected with a gentleMACS dissociator (Miltenyi, Bergisch Gladbach, Germany) and then treated with collagenase I and DNase (Roche, Basel, Switzerland) for 45 min at 37°c. The cells were filtered and stained with antibodies. The following antibodies were used in this study: anti-mouse CD11c, anti-mouse/human CD11b, anti-mouse Ly6G, anti-mouse F4/80, anti-mouse CD69, anti-mouse CD8a, anti-mouse CD4 anti-mouse Ly6C, anti-mouse I-A/I-E, anti-mouse CD25, anti-mouse CD40, Fixable Viability Dye eFluor® 450 (BioLegend, CA, USA). Cell apoptosis, CFSE, H2DCFDA, and MitoTracker Green staining were measured under the manufacturer protocols (Life Technologies, USA). BD FACS Canto flow cytometer (BD, NJ, USA) and FlowJo software were used for flow cytometry analysis (TreeStar, OR, USA). Antibodies used in this set of experiments are listed in Supplementary Table 1.

### In vitro MDSC Generation

MDSCs were generated as previously reported^35^. Briefly, bone marrow nucleated cells were cultured in DMEM supplied with 2mM L-glutamine, 1.5g/L sodium bicarbonate, 10mM HEPES, 1.0mM sodium pyruvate (all from Gibco, MA, USA), 10% fetal bovine serum (Hyclone, MA, USA), 40ng/ml rmGM-CSF and 40ng/ml rmIL-6 (BioLegend), with or without 50% E.G7-OVA supernatant. The floating cells were collected on day 4 and enriched by anti-mouse Gr-1 Microbeads (Miltenyi). The cells with purity > 90% were used. In some experiments, the cells were stained with anti-mouse Ly6C, anti-mouse Ly6G, and anti-mouse/human CD11b (Biolegend) and sorted with BD FACS Aria II cell sorter (BD).

### T-cell proliferation assay

MDSCs were prepared as described above. T cells were separated from spleens of C57BL/6 mice and enriched using the PanT Cell Isolation Kit II (Miltenyi). 2×10^5^ T cells labeled with CFSE (MitoSciences, OR, USA) and 1×10^5^ MDSC were seeded into 96-well plates (Corning, NY, USA). Mouse T-activator CD3/CD28 beads (ThermoFisher Scientific, MA, USA) were added to the culture media. The cells were cultured in a humidified incubator with 5% CO2 for 72h and analyzed using a BD FACS Canto flow cytometer (BD,) and FlowJo software (TreeStar, OR, USA).

### ELISpot assay

Single-cell suspensions were generated from the spleens of WT and *Camkk2^-/-^* mice 17 days after E.G7-OVA injection. Splenocytes (2×10^5^/well) were seeded into 96-well MultiScreen Immubilon-P Filtration Plates (Milllipore, USA) coated with anti-mouse IFN-γ antibody (Affymetric eBioscience, USA). OVA protein was added at 100μ/ml. Cells were cultured in a humidified incubator with 5% CO2 for 48 h. The reactions were processed according to the protocol instructions. CTL-ImmunoSpot® S6 FluoroSpot Line (Cellular Technology Ltd, OH, USA) and ImmunoSpot software (Cellular Technology Ltd, OH, USA) was used for analysis.

### Immunoblot

Immunoblots were prepared as published^24^. The following primary antibodies were used in this study: purified mouse anti-CaM kinase kinase (BD Biosciences, CA, USA), anti-phospho-AMPK alpha (T172 40H9) rabbit mAb, anti-AMPK alpha (F6) mouse mAb, anti-phopho-Stat3 (Tyr705 D3A7) rabbit mAb, anti-Stat3 (124H6) mouse mAb, anti-β actin mouse mAb (all from Cell Signaling, CA, USA). Secondary antibodies were anti-mouse IgG Alexa Fluor 680 (Invitrogen, CA, USA) or anti-rabbit IgG IRDye800 conjugated (Rockland Immunochemicals, PA, USA). The image was taken using Odyssey CLx (LI-COR, NE, USA) and analyzed using ImageStudio (LI-COR, NE, USA). Antibodies used in this set of experiments are listed in Supplementary Table 2.

### Real-Time Quantitative RT-PCR Assay

RNA was extracted using the RNeasy Mini Kit (Qiagen, Hilden, Germany). Quantitative PCR was performed using iQ SYBR Green Supermix (Bio-Rad, CA, USA) with respective primers and cDNA. The assay was run on the CFX96 Real-Time System (Bio-Rad, CA, USA). The primer sequences are listed in Supplementary Table 3.

### Cell Metabolic Assay

OCR and ECAR were measured with the Seahorse XF24 extracellular flux analyzer (Agilent Technologies. CA, USA). Cells were attached to culture plates using Cell-Tak (BD Bioscience). OCR and ECAR were measured in un-buffered DMEM supplied with 4.5g/L D-glucose and 10mM L-glutamine (Gibco, MA, USA). OCR and ECAR values were normalized to cell numbers. Data were analyzed using Seahorse Wave (Agilent Technologies. CA, USA).

### Statistical Analysis

All statistical analyses were performed using Prism GraphPad (GraphPad Software, CA, USA) and Excel (Microsoft, WA, USA). For survival studies, the log-rank Mantel-Cox test was used. For tumor size and bioluminescent measurement, student *t*-test and multi-way ANOVA tests were used. For flow cytometry, real-time quantitative RT-PCR assay, and cytokine detection analysis, a two-tail student *t*-test was used. The level of significance was set at *P < 0.05*. Bar graphs represent mean ± SEM.

## Results

### Deletion of *Camkk2* in the host inhibits lymphoma cells growth

Outside the brain, we originally found CaMKK2 selectively expressed in myeloid cells and macrophages where this kinase regulates the inflammatory response triggered by bacterial endotoxin^25^. More recently, we have demonstrated that CaMKK2 expressed in myeloid cells is an important regulator of the immunosuppressive breast cancer microenvironment and blockade of this kinase in the host cells inhibits tumor growth^26^. Based on these results, we hypothesize that, besides breast cancer, CaMKK2 expressed in the host cells might have a more general function in the mechanism regulating the anti-tumor immune response. Consequently, we used E.G7-OVA tumor cells, a well-established derivative model of the EL-4 murine lymphoma cells, which has been modified to express an ectopic T cell-recognizable ovalbumin peptide^36^. The E.G7-OVA cells were further engineered to express luciferin as a trackable marker so bioluminescent imaging was combined with caliper measurements to monitor tumor development. E.G7-OVA cells were thus injected subcutaneously (SC) into the right flank of wild type (WT) and *Camkk2^-/-^* mice and tumors monitored by bioluminescence and caliper. In WT mice injected with 1×10^4^ E.G7-OVA cells, the tumors were detectable by bioluminescence by day 7 and gradually increased over the following two weeks (**Figs. 1A and S1A**). Although tumors successfully engrafted in *Camkk2^-/-^* mice, the bioluminescent signal gradually decreased after 21 days and eventually disappeared after 32 days. To further corroborate this finding, the experiment was replicated using an increased number of tumor cells (1×10^5^). However, even in the conditions of increased tumor cell input, we found that deletion of CaMKK2 in the host cells effectively suppressed syngeneic tumor cell growth (**Fig. 1B**).

**Figure 1.**
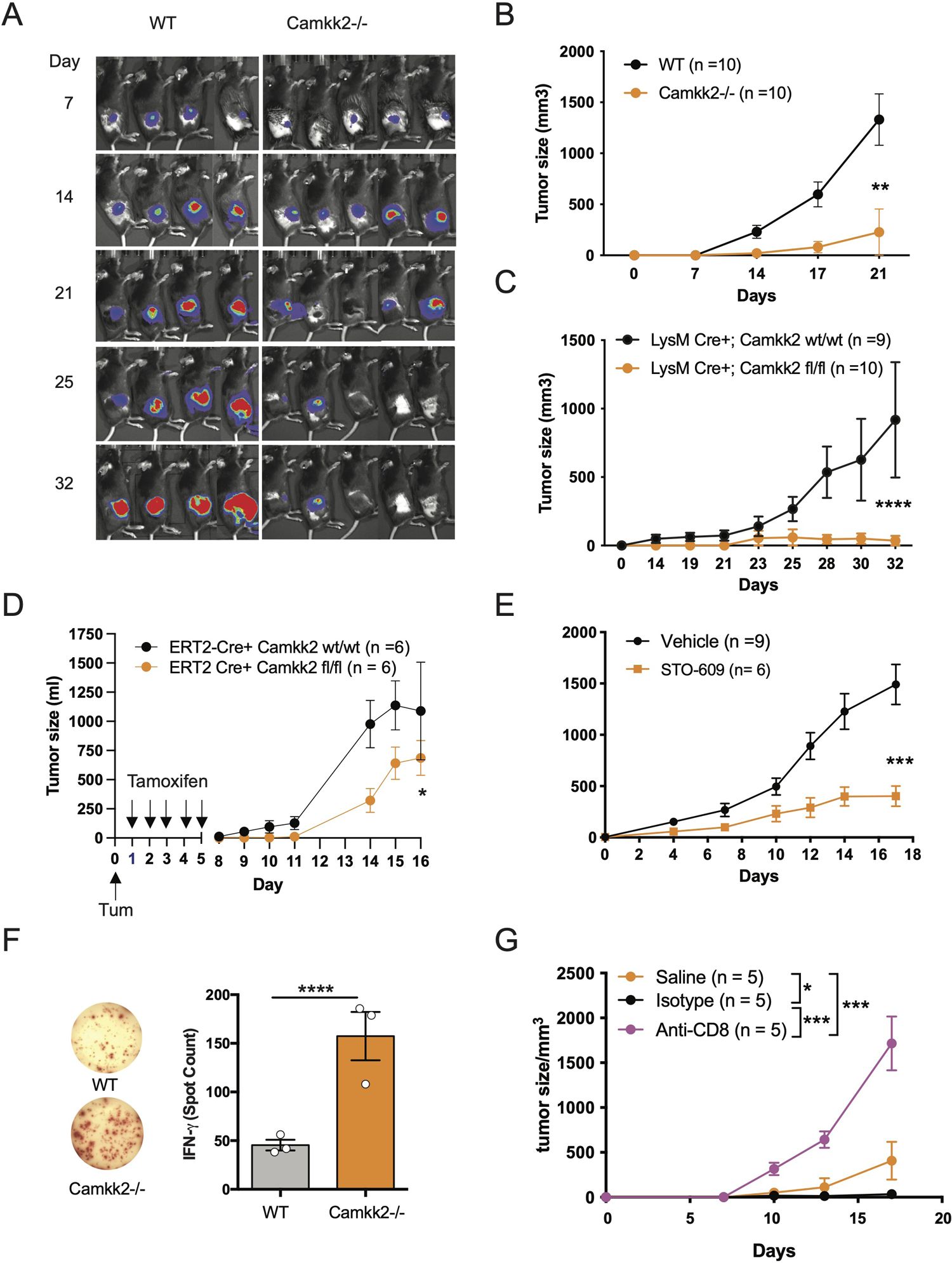
Deletion of *Camkk2* in the host inhibits lymphoma cells growth. WT and *Camkk2^-/-^* mice were injected with E.G7-OVA s.c. in the flank. The tumors were measured every 3-4 days. (A) Bioluminescent imaging of tumors growing in WT and *Camkk2^-/-^* mice (1×10^4^ cells/mouse). (B) Tumors size in WT and *Camkk2^-/-^* mice (1×10^5^ cells/mouse). N = 10 mice/group. Repeated twice. (C) Tumors size in LysM-Cre+; CaMKK2^wt/wt^ and LysM-Cre+; CaMKK2^fl/fl^ mice (5×10^5^ cells/mouse). N = 9 LysM-Cre+; CaMKK2^wt/wt^ and 10 LysM-Cre+; CaMKK2^fl/fl^ mice. Combined from two experiments. (D) ERT2-Cre Camkk2^loxp^ mice were injected with EL4 cells s.c. in the flank and treated with tamoxifen (20 mg/kg, daily, oral gavage). Tumor growth was monitored by caliper. (E) WT mice were injected with EL4 cells s.c. in the flank and treated with STO-609 (40 μmoles/Kg, IP at 48 hours interval, total 7-doses). (F) WT and *Camkk2^-/-^* mice were injected with 1×10^5^ E.G7-OVA s.c. in the flank. After 17 days the spleens were removed, and anti-OVA T-cell response was assessed in splenocytes. N=3, repeated twice. (G) E.G7-OVA tumor growth in *Camkk2^-/-^* mice treated with saline, anti-CD8+ or isotype control antibodies. N=4-5. Repeated twice. *, **, *** and **** refer to p< 0.05, 0.01, 0.005, and 0.001 respectively.

We next examined the cell types expressing *Camkk2* in the tumor microenvironment and spleen of lymphoma-bearing mice by engrafting E.G7-OVA cells into the flank of [Tg(*Camkk2*-EGFP)C57BL/6J] *Camkk2*-reporter mice^27^. After 3 weeks the tumors and spleens were removed and embedded in paraffin for histology or enzymatically digested to prepare a single-cell suspension. We then used immunostaining and flow cytometry analyses to examine immune cell subsets^37^. There were a significant number of stromal cells associated with E.G7-OVA tumors expressing EGFP reporter protein (**Fig. S1B**). Flow cytometry analysis confirmed this result and showed robust *Camkk2*-reporter activity in tumor-associated CD11b^+^ myeloid cells, but minimal activity in CD11b^-^ non-myeloid cells (**Fig. S1C, left panel**). To identify myeloid cell subsets, CD11b^+^ cells were further sub-gated using Ly6C, Ly6G, F4/80, and MHC II markers (**Figs. S1C-D**). The *Camkk2* promoter was highly active in the Ly6C^high^/Ly6G^dim^ subset (M-MDSC) and CD11b^+^, Ly6C^dim^, Ly6G^dim^, F4/80^+^ tumor-associated macrophage-like cells (**Figs. S1C, lower left panel, DN gate**). Conversely, we observed low levels of EGFP in the Ly6C^dim^/Ly6G^high^ subset (G-MDSC) and Ly6C^neg^/Ly6G^neg^/F4/80^dim/neg^/MHC-II^dim/neg^ cells (**Fig. S1C**). We next evaluated the expression of EGPF in the spleen from E.G7-OVA-bearing mice and control groups. M-MDSC and DN splenocytes expressed the highest levels of EGFP reporter regardless of tumor status. However, the EGFP reporter was expressed at low levels in G-MDSC and barely detectable in CD11b^-^ cells (**Fig. S1D**). These results indicate the *Camkk2* promoter is highly active in tumor-associated myeloid cells and in myeloid subsets in the spleen of tumor-bearing mice.

We originally reported that CaMKK2 is significantly expressed in macrophages associated with the breast cancer microenvironment^26^. Here we extend this finding to the lymphoma microenvironment, and more importantly, we identified spleen and tumor residing MDSCs as relevant cell types expressing this kinase. To determine the functional relevance of CaMKK2 expressed in myeloid cells in our model of hematological malignancy, we used LyMCre^+^ *Camkk2*^loxP^ mice in which *Camkk2* is selectively deleted in the myeloid compartment^26^. LyMCre^+^ *Camkk2*^fl/fl^ and LyMCre^+^ *Camkk2*^wt/wt^ littermates were thus challenged with E.G7-OVA cells subcutaneously. Consistent with our observations in the *Camkk2^-/-^* global knockout model, the E.G7-OVA tumors were significantly smaller in mice genetically deleted of *Camkk2* in the myeloid cells compared to control littermates (**Fig. 1C**). To determine the potential translational implications of this finding, we next leveraged our model of inducible *Camkk2* knockout mice. We first demonstrated that treatment with tamoxifen was able to induce a significant downregulation of *Camkk2* mRNA expression in the relevant myeloid cell population. Mice were treated with tamoxifen and then *Camkk2* RNA expression was determined by QT-PCR in peritoneal lavage macrophages isolated from ERT2-Cre^+^ Camkk2^fl/fl^ and ERT2-Cre^+^ Camkk2^fl/fl^. The results from this experiment indicate that, following tamoxifen treatment, the expression of *Camkk2* RNA was significantly lower in ERT2-Cre^+^ Camkk2^fl/fl^ compared to control littermates (**Fig. S1E**). EL-4 cells were next SC inoculated in ERT2-Cre^+^ Camkk2^fl/fl^ mice and their control littermates (ERT2-Cre^+^ Camkk2^wt/wt^) and mice were then treated with tamoxifen. The results of these studies indicated that the acute deletion of *Camkk2* in the host cells was sufficient to induce significant inhibition of the lymphoma cells growth (**Fig. 1D**). Lastly, we evaluated the efficacy of STO-609, a small molecule pharmacological CaMKK2 inhibitor^38^, to suppress the growth of EL-4 lymphoma cells. Mice were injected with EL-4 cells and 24 hours later were treated with STO-609 for 2 weeks. Similar to findings observed in genetically modified models of *Camkk2* deletion (**Figs. 1A-D**), tumors growing in mice treated with STO-609 were significantly smaller compared to mice receiving vehicle alone (**Fig. 1E**). Although this pharmacological approach cannot distinguish the intrinsic effect of STO-609 on tumor cells from those on stroma cells, it provides further evidence on the potential translational impact of targeting CaMKK2 to restrain lymphoma growth.

### Deletion of *Camkk2* in the host enhances anti-tumor T-cell mediated immunity

CD8^+^ T cells have an important role in the mechanism of tumor rejection and consequently, an increased number of CD8^+^ tumor-infiltrating lymphocytes (TIL) has been associated with a good prognosis in several tumor types^39^. Moreover, we have previously shown that deletion of CaMKK2 in the host restrains mammary cancer growth, and this phenomenon was associated with increased accumulation of T cells in the tumor microenvironment and prevented by depletion of CD8^+^ T cells^26^. To determine whether a similar mechanism is also present in the lymphoma tumor model, we first analyzed the percentage and phenotype of TILs in EG-7-OVA tumors in WT and *Camkk2^-/-^* mice. To this end, E.G7-OVA cells were injected into the flank and the TIL phenotype was then analyzed by FACS in tumors removed 14-17 days later. The data indicated comparable percentages of both CD4^+^ and CD8^+^ T cells in tumors removed from *Camkk2^-/-^* and WT mice (**Figs. S2A-B**). However, the percentage of CD8+ GranzymeB^+^ PD1^-^ TIL, a phenotype associated with non-exhausted effector T cells, was increased in E.G7-OVA tumor removed from *Camkk2^-/ -^* compared to WT mice (**Fig. S2B**). Of note, changes in the TIL repertoire observed in lymphoma models resemble those originally described in mammary tumors growing in *Camkk2^-/ -^* compared to WT mice^26^, indicating that in these two models of tumors deletion of *Camkk2* in the host trigger a similar anti-tumor effector mechanism.

We next leveraged the EG7-OVA lymphoma model to determine the ability of WT and *Camkk2^-/ -^* mice to develop a specific T cell-mediated response toward the OVA antigen, which is selectively expressed by E.G7-OVA lymphoma cells. Fresh splenocytes were isolated from E.G7-OVA-bearing mice and then cultured *in vitro* in the presence of OVA protein. The cells were then analyzed for IFN-γ production by ELISpot. We found three-fold more IFN-γ spots in splenocytes isolated from *Camkk2^-/ -^* compared to WT mice, which suggests the decrease of MDSC in *Camkk2^-/-^* mice unleashes an anti-tumor immune response that restrains tumor growth (**Fig. 1F**). Lastly, we determined the role of CD8+ T-cell-mediated immune response in the lymphoma growth inhibition mechanism. To this end, *Camkk2^-/-^* mice were treated with anti-CD8 depleting antibody or isotype control and subsequently challenged with E.G7-OVA cells (**Fig. S2C)**. Tumor growth was suppressed in *Camkk2^-/-^* mice that received no treatment or isotype IgG control, while the treatment with anti-CD8 depleting antibody reversed the protective effects of *Camkk2* deletion, enabling the growth of E.G7-OVA (**Fig. 1G**). These findings indicated that the loss of CaMKK2 in the host cells enhances the anti-tumor T cell response, and in turn suppressess lymphoma cells to grow in *Camkk2^-/-^* mice.

### Deletion of *Camkk2* in the host prevents tumor-induced MDSC expansion

The accumulation of MDSCs in lymphoid tissues and blood of tumor-bearing individuals is an important immunosuppressive phenomenon associated with poor prognosis in preclinical tumor models and humans^5^. We thus hypothesized that CaMKK2 regulates myeloid cell responsiveness to tumor-derived factors, and inhibition of this kinase in host cells prevents the tumor-induced MDSC expansion. To test this hypothesis, we determined the effect of *Camkk2* deletion on MDSCs accumulating in the spleen of normal and tumor-bearing mice. In normal conditions (CTR), WT and *Camkk2^-/-^* mice had comparable CD11b^+^ and Ly6C^high^ Ly6G^dim^ percentage in the spleens, while *Camkk2^-/-^* mice had a slight increase of Ly6G^dim^ Ly6G^high^ compared to WT (**Figs. 2A-B**). Regardless of the genotype, a significant increase in the percentage of MDSCs was detected in EG7-OVA-bearing (TB) mice compared to the control groups (**Fig. 2B**). Interestingly, less MDSC accumulated in the spleens and in the tumors of *Camkk2^-/-^* compared to WT mice (**Figs. 2B-C**).

**Figure 2.**
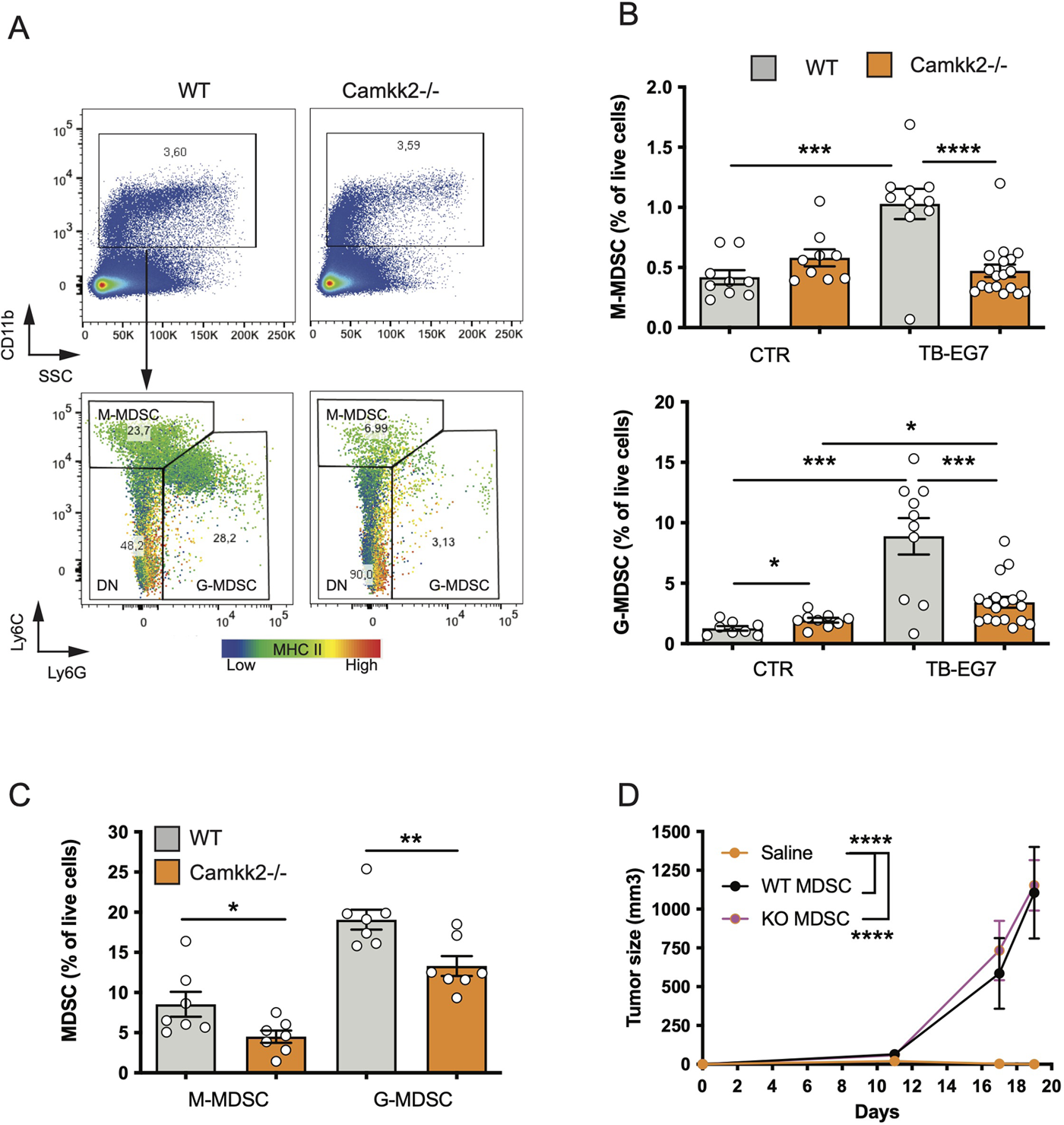
Deletion of *Camkk2* in the host prevents tumor-induced MDSC expansion. WT and *Camkk2^-/-^* mice were injected with E.G7-OVA s.c. in the flank (1×10^5^ cells/mouse). Spleens and tumors were removed, and single cell suspensions were stained for myeloid markers and analyzed by FACS. Groups of WT and *Camkk2^-/-^* mice not injected with tumor cells were used as control. (A) Gating strategy and representative plots of splenocytes isolated from tumor-bearing WT and *Camkk2^-/-^* mice 17-21 days after injection.The heatmap indicates MHC II intensity in B and Ly6C intensity in C, respectively. (B) Percentages of M-MDSC, G-MDSC subsets in the spleen of normal and tumor-bearing mice (CTR and TB-EG7, respectively). (C) MDSCs in tumors removed from WT and *Camkk2^-/-^* mice. (D) 1×10^4^ E.G7-OVA were injected s.c. in the flank of *Camkk2^-/-^* mice. At the same time, mice were injected IV with Gr1+ MDSC isolated from the spleen of tumor-bearing WT or *Camkk2^-/-^* mice. The vehicle group received an equal volume of saline only. N=5 mice/group. Repeated twice. *, **, *** and **** refer to p< 0.05, 0.01, 0.005, and 0.001 respectively.

To evaluate whether the failure to expand MDSCs has an important causative role in the mechanism of tumor suppression, MDSCs were isolated from the spleens of E.G7-OVA-bearing WT and *Camkk2^-/ -^* then adoptively transferred into *Camkk2^-/-^* mice via tail vein injection. *Camkk2^-/-^* mice receiving sham tail vein injection were also included as a control group. EG7-OVA cells were then inoculated in the flank of CTR and MDSC-treated mice. EG-7-OVA cells failed to grow robustly in the *Camkk2^-/-^* control group (**Fig. 2D**), while tumors developed at an equal rate in *Camkk2^-/ -^* mice that received either WT or *Camkk2^-/ -^* MDSCs (**Fig. 2D**). Restoring the number of MDSC was thus sufficient to revert the protective effect of *Camkk2* deletion on tumor growth, supporting the hypothesis that the failure to expand MDSC has an important causative role in the mechanism of tumor suppression in *Camkk2^-/-^* mice. Further, this data also indicated that WT and *Camkk2^-/-^* MDSCs have a comparable immunosuppressive activity, at the cellular level.

### Deletion of *Camkk2* impairs MDSC expansion *in vitro*

To establish the cell-intrinsic function of CaMKK2 in MDSCs, we leveraged well-characterized models of *in vitro* bone marrow-derived MDSC (BM-MDSC)^40, 41^. We first determined whether bone marrow cells isolated from WT and *Camkk2^-/-^* mice have a comparable ability to generate MDSC. To this end, bone marrow cells were collected from WT and *Camkk2^-/-^* femurs and cultured with GM-CSF/IL-6 in the presence or absence of E.G7-OVA-conditioned medium (TCM). After 4 days, non-adherent cells were harvested and analyzed for phenotype and function. The expression of CaMKK2 at the mRNA and protein levels was assessed using qPCR and immunoblot, respectively (**Figs. S3A-B**), while phenotype and yield of MDSC subsets were evaluated by flow cytometry (**Figs. 3A-B**). The results of these experiments indicated that CaMKK2 was expressed at detectable RNA and protein levels by *in vitro* generated MDSC and were not detected in *Camkk2*^-/-^ MDSC (**Figs. S3A-B**). Less MDSC were harvested from bone marrow *Camkk2*^-/-^ cultures compared to WT (**Figs. 3A-C**). Of note, this phenomenon was associated with an increased percentage of cells showing a phenotype of more differentiated myeloid cells (CD11b+, Ly6C^neg^, Ly6G^neg^), which included more differentiated populations of non-adherent myeloid cells (e.g., dendritic cells; **Fig. 3B**). To determine whether the effects of *Camkk2* were reproducible in other experimental settings, BM cells from WT and *Camkk2*^-/-^ mice were cultured with GM-CSF in the presence or absence of IL-18, which has been identified as an important driver of MDSC-mediated immunosuppression^42, 43^. In addition, MDSC from both genotypes were also generated in the presence or absence of TCM derived from several breast cancer cell lines, including EO771, 4T1, MET1, and A7C11. Consistent with the results from GM-CSF/IL-6 and EG7-OVA-conditioned medium, in all these experimental conditions, fewer MDSCs were generated from *Camkk2^-/-^* BM cells compared to WT (**Fig. 3D**). Lastly, WT and *Camkk2*^-/-^ BM cells were cultured with GM-CSF in the presence or absence of STO-609 (2 µM) or vehicle. After 4 days cells were harvested, and the phenotype was determined by flow cytometry. With results from BM cells isolated from WT and *Camkk2^-/-^* mice, the presence of STO-609 in the differentiation medium impaired the ability of WT BM cells to generate MDSCs (Fig. 3E). In contrast, STO-609 slightly affected the yield of M-MDSC and exerted no significant changes on G-MDSC generated from *Camkk2^-/-^* BM cells (**Fig. 3E**).

**Figure 3.**
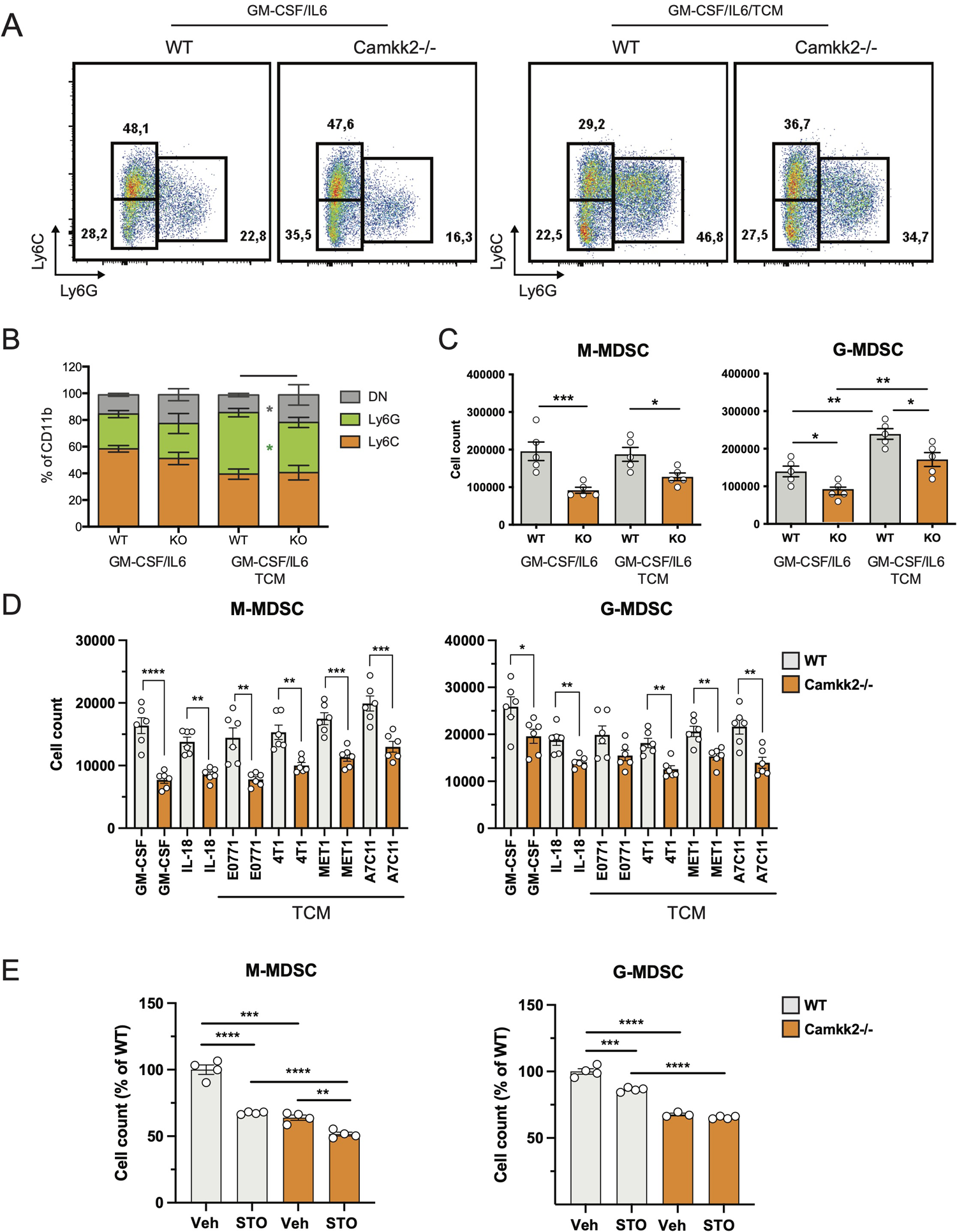
Deletion of *Camkk2* impairs MDSC expansion *in vitro.* Bone marrow nucleated cells from WT and *Camkk2^-/-^* mice were cultured with cytokines in the absence or presence of E.G7-OVA tumor conditioned medium (30% v/v). After 4 days, cells were collected, stained, and analyzed by flow cytometry. (A) Gating strategy to identify MDSCs and representative dot plots. (B-C) Percentage and absolute number of MDSCs. (D). Yield of MDSC generated with GM-CSF in the presence of the cytokines and mammary tumor cells-TCM (E) Yield of MDSC generated in the presence of STO-609 (2mM). *, **, *** and **** refer to p<0.05, 0.01, 0.005, and 0.001, respectively.

The impact of *Camkk2*-deficiency on the immunosuppressive functions of MDSC was evaluated next. For these studies, MDSC were generated from BM of WT and *Camkk2^-/-^* MDSC. After 4 days, MDSC were collected and co-cultured with CFSE-labeled T cells with or without anti-CD3/CD28 beads. Regardless of genotype, MDSC exerted significant suppressive effects on T cells, inhibiting proliferation, CD25, and CD69 upregulation induced by CD3/CD28 stimulation (**Figs. S3C-D**). Cumulatively, these data confirmed the *in vivo* tumor suppression results, indicating that the deletion of *Camkk2* impacts MDSC expansion, exerting only a negligible effect on MDSC immunosuppressive function.

### Deletion of *Camkk2* impairs survival and enhances terminal differentiation of MDSC

Multiple mechanisms could account for the decreased expansion of *Camkk2^-/-^* MDSC such as increased cell death or enhanced differentiation into mature myeloid cells (e.g. macrophages and dendritic cells). We first investigated whether the deletion of *Camkk2^-/-^* increased the susceptibility of MDSC to apoptosis. Bone marrow cells from WT and *Camkk2^-/-^* were cultured in the presence of GM-CSF and EG7-OVA-CM. After 4 days, MDSC were harvested and apoptotic cells were identified by flow cytometry. As expected, less M-MDSC and G-MDSC were recovered from *Camkk2^-/-^* BM cultures compared to WT (**Figs. 4A-B**). Consistent with this result, higher percentages of early and late apoptotic cells (AnnexinV^+^ 7ADD^-^ and AnnexinV^+^ 7-AAD^+^, respectively) were also detected in *Camkk2^-/-^* MDSC subsets compared to WT (**Fig. 4C**). This finding prompted us to evaluate the capability of *Camkk2^-/-^* MDSC to survive during oxidative stress, a well-known inducer of cell death^44^. MDSC cells were then exposed to hydrogen peroxide (H2O2)-induced oxidative injury and the percentage of apoptotic cells was monitored over time by flow cytometry. A progressive increase in either M-MDSC and G-MDSC apoptotic cells was observed following acute exposure to H2O2. Of note, the sensitivity of MDSC generated from *Camkk2^-/-^* BM cells to the hydrogen peroxide oxidative stress- was significantly higher compared to WT (**Fig. 4D**).

**Figure 4.**
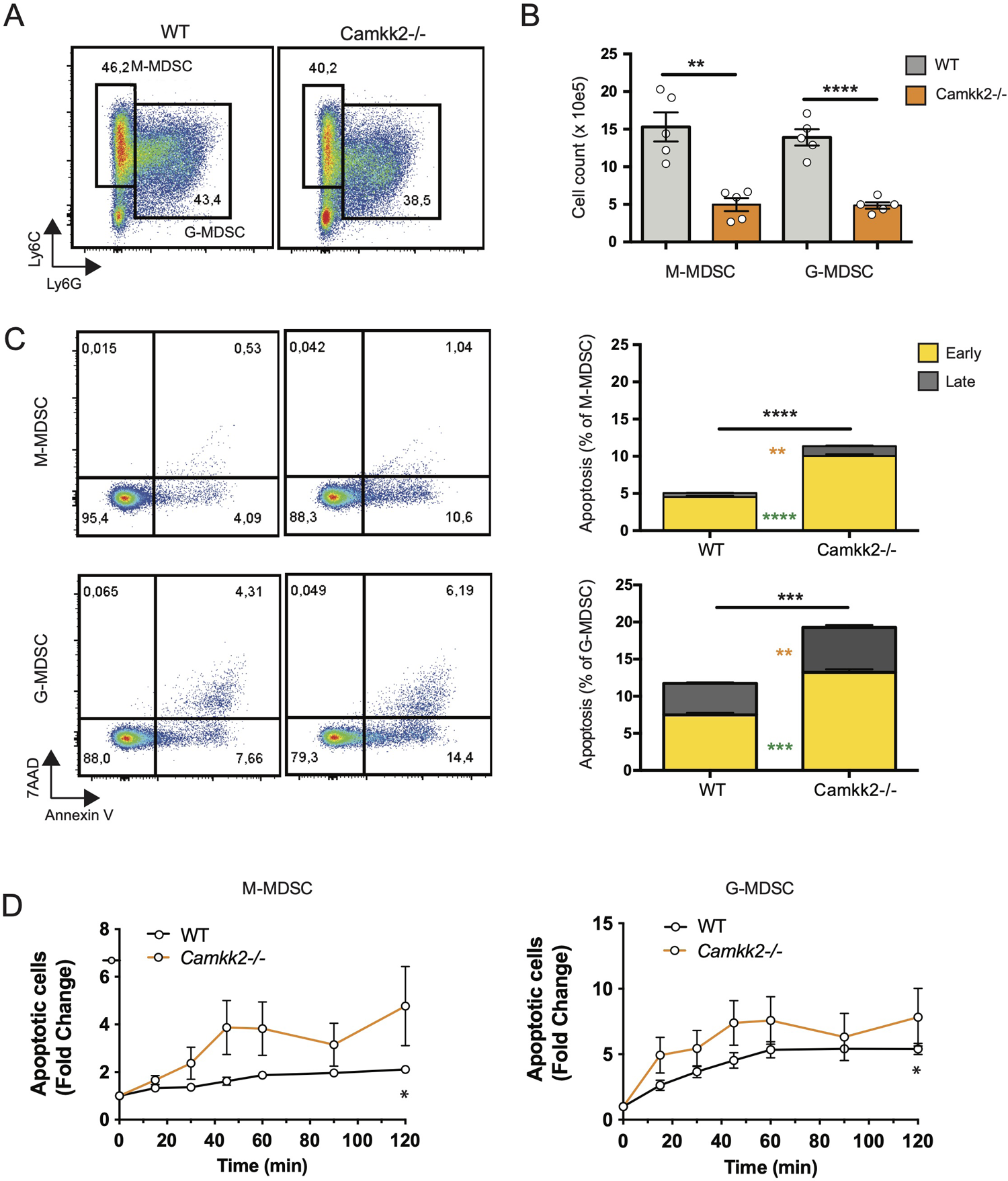
Deletion of *Camkk2* impairs the survival of MDSC. Bone marrow cells from WT and *Camkk2^-/-^* mice were cultured with GM-CSF/IL6 cytokines in the absence or presence of E.G7-OVA tumor conditioned medium (30% v/v). After 4 days, cells were collected, stained, and analyzed by flow cytometry. (A) Gating strategy and representative FACS profiles. (B) Yield of WT and *Camkk2^-/-^* M-MDSC, G-MDSC, and DC. (C) Percentage of early and late apoptotic MDSC. N = 5 biological replicates/group. (D) Bone marrow nucleated cells from WT and *Camkk2^-/-^* mice were cultured with GM-CSF/IL6 cytokines in the absence or presence of E.G7-OVA tumor conditioned medium (30% v/v). MDSC were collected after 48 hours of culture and exposed to 1μ H2O2. Apoptotic cells were identified by flow cytometry at the indicated time points. *, **, ***, and **** refer to p<0.05, p< 0.01, p<0.001, p<0.0001, respectively. T-test and two-ways ANOVA were used to calculate p-values in panels B and D, respectively.

Data combined from multiple experiments indicated that, alongside a decreased yield of MDSC, more mature myeloid cells showing the typical phenotype of DC (Ly6^-^, Ly6G^-^, CD11c^+^, MHC II^high^) were generated from *Camkk2^-/-^* BM cells cultures compared to WT (**Figs. 5A-B**). Further, adherent cells recovered from WT BM cultures exhibited a typical M2-like morphology with significant elongation and branching while the cells recovered from *Camkk2^-/-^* cultures displayed a round shape M1-like morphology (**Fig. S4A**). FACS and qPCR analyses corroborated this data, showing higher MHC II expression and lower *ArgI* and *Chi3l3* mRNAs levels in the adherent cells recovered from *Camkk2^-/-^* cultures compared to WT cells (**Figs. S4B-C**). These findings indicated that, in addition to its effects on apoptosis, CaMKK2 functions to fine-tuning the terminal differentiation process of more mature MDSC. To directly evaluated this hypothesis, *in vitro* generated M-MDSC (Ly6C^high^ Ly6G^dim^ CD11c^neg^ MHC II^neg^) and G-MDSC (Ly6C^dim^ Ly6G^high^ CD11c^neg^ MHC II^neg^) were sorted by FACS and cultured with G-MCSF/IL6 in the presence of E.G7-OVA-TCM for additional 48 hours. Non-adherent cells were then harvested and analyzed by flow cytometry. As expected, the terminal differentiation of M-MDSC was associated with a consistent downregulation of Ly6C expression, leading to a remarkable accumulation of LyC^dim^/Ly6G^dim^ DN cells (**Figs. 5C top panels, orange dots, 5D**). Interestingly, in DN gate, more cells expressing the typical phenotype of dendritic cells (CD11c^+^ MHC II^high^) were generated from *Camkk2^-/-^* purified M-MDSC compared to WT (**Figs. 5C lower panels, orange dots, and 5E**). Regardless of genotype, the majority of cells recovered from G-MDSC cultures still expressed detectable levels of Ly6G and only a negligible number of them acquired DN/DC phenotypes (**Figs. 5C, green dots)**. Collectively, these findings indicated that deletion of *Camkk2* accelerates the terminal differentiation of M-MDSC cells toward more mature myeloid cell subsets (e.g., M1 macrophages and dendritic cells) and this mechanism, in conjunction with increased apoptosis would account for the insufficient expansion of *Camkk2^-/-^* MDSC.

**Figure 5.**
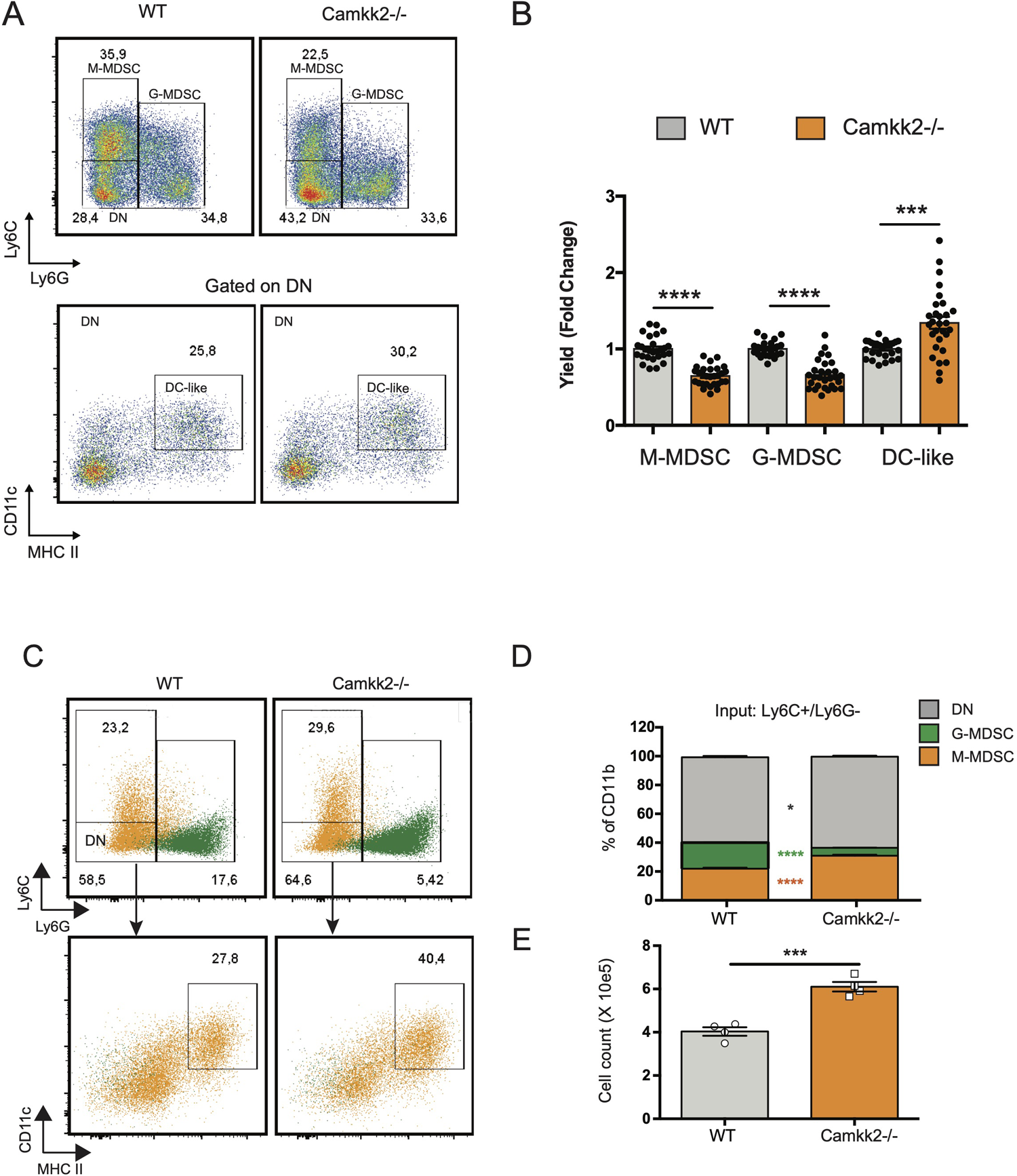
Deletion of *Camkk2* accelerates the terminal differentiation of MDSCs. Bone marrow cells from WT and *Camkk2^-/-^* mice were cultured with GM-CSF/IL6 cytokines in the absence or presence of E.G7-OVA tumor conditioned medium (30% v/v). After 4 days, cells were collected, stained, and analyzed by flow cytometry. (A) Gating strategy and representative FACS profiles. (B) Yield of M-MDSC, G-MDSC and DC. Bar graph showing the cumulative results from 6 independent experiments and fold changes (FC) have been calculated on the yield of WT MDSC. (C-D) Bone marrow cells from WT and *Camkk2^-/-^* mice were cultured with GM-CSF/IL6 cytokines in the absence or presence of E.G7-OVA tumor conditioned medium (30% v/v). After 4 days, non-adherent cells were collected and sorted by flow cytometry. Purified M-MDSC and G-MDSC were then cultured in the presence of GM-CSF/IL-6 and 30% of EG7 CM for additional 48 hours and analyzed by flow cytometry. (C) Gating strategy to identify M-MDSC, G-MDSC, and DC in M-MDSC and G-MDSC cultures. Overlapping dot plots refer to the FACS profiles of cells collected from M-MDSC and G-MDSC (shown in orange and green, respectively). (D) Percentage of M-MDSC, G-MDSC, and DN subsets (upper). (E) Yield of Ly6C-Ly6G-CD11c^+^ MHC II^+^ cells (namely DC). *, ** and *** refer to p<0.05, p< 0.01 and 0.005, respectively.

### *Camkk2* deficiency increases reactive oxygen species (ROS) accumulation and impairs mitochondria functions in MDSC

ROS are important regulators of multiple cellular processes including cell death, proliferation, and differentiation^44, 45^. Physiological levels of ROS are required for MDSC generation^46^, while genetic alterations causing abnormal accumulation of ROS trigger MDSC apoptosis, which prevents tumor-induced MDSC expansion^47, 48^. Increasing evidence documents a network of relevant bidirectional interactions among ROS and calcium signaling^49–51^, and CaMKK2 has been recently identified as an important regulator of ROS accumulation and ferroptosis in tumor cells^52^. These findings prompted us to determine whether deletion of *Camkk2* interferes with the mechanism of ROS accumulation in MDSCs. To test this, MDSCs were generated from BM cells of WT and *Camkk2^-/-^* mice, and ROS accumulation was evaluated by flow cytometry (**Fig. S5A**). The results of these experiments indicated that higher ROS levels accumulated in *Camkk2^-/-^* MDSC compared to WT (**Fig. 6A**). We next evaluated whether deletion of *Camkk2* interferes with the expression of genes involved in ROS production (NADPH Oxidase 1 and 2; *Nox1* and *Nox2,* respectively) or in the antioxidant response (Catalase and nuclear factor erythroid 2-related factor-2, *Cat* and *Nrf2*, respectively). Genes positively regulating ROS accumulation (*Nox1* and *Nox2*) were expressed at comparable (*Nox1*) or lower levels (*Nox2*) in *Camkk2^-/-^* MDSC compared to WT (**Fig. S5B**). Moreover, lower levels of *Cat* and *Nfr2* mRNAs were expressed in *Camkk2^-/-^* MDSC compared to WT (**Figs. 6B and S5B**). These findings, alongside the higher susceptibility of *Camkk2*^-/-^ MDSC to oxidative stress-induced apoptosis (**Fig. 4D**), indicated that CaMKK2 could function by regulating the antioxidant response in MDSC. To evaluate the functional role of ROS, MDSC were generated from BM of WT and *Camkk2^-/-^* mice in the presence or absence of hydrogen peroxide^44^ or butylated hydroxyanisole (BHA), which is a known scavenger of ROS^53^. The presence of hydrogen peroxide in the culture medium impaired the production of WT MDSC, with no significant effects on *Camkk2^-/-^* M-MDSC (**Fig. S5C upper**). Further, regardless of the genotype, the yield of G-MDSC and DC was negatively regulated by prolonged exposure to H2O2 (**Fig. S5C upper**). On the contrary, more MDSC and less DC were recovered from the culture of WT bone marrow cells treated with BHA (**Fig. S5C, lower)**. However, only minor effects were induced by this ROS scavenger on *Camkk2^-/-^* MDSC and DC (**Fig. S5C, lower)**. We lastly tested the ability of MitoTempo, a specific scavenger of mitochondrial superoxide^54^ to interfere with the generation of MDSC. As predicted by our hypothesis, more MDSC were recovered from WT MDSC treated with this ROS scavenger (**Figs. 6C-D**). Interestingly, the targeting of the mitochondrial source of ROS with MitoTempo also increased the yield of *Camkk2^-/-^* M-MDSC and G-MDSC cells (**Figs. 6C-D**). On the other hand, treatment with MitoTempo was associated with a decreased yield of WT and *Camkk2^-/-^* DC (**Fig. S5D**).

**Figure 6.**
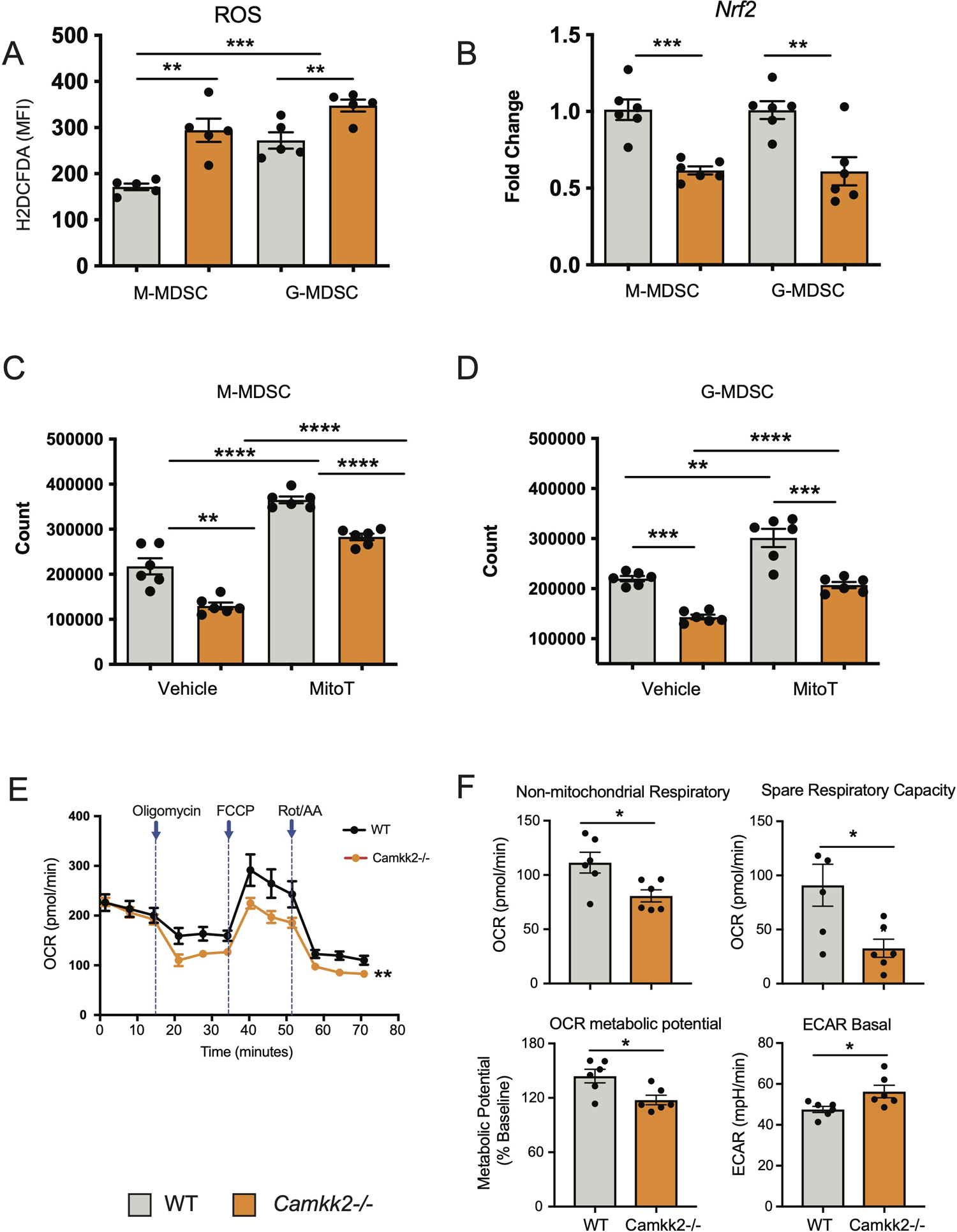
*Camkk2* deficiency increases reactive oxygen species (ROS) accumulation and impairs mitochondria functions in MDSC. MDSCs were generated from bone marrow nucleated cells of WT and *Camkk2^-/-^* mice femurs. (A) MDSCs were identified according to the gating strategy described previously and ROS levels were evaluated by flow cytometry using H2DCFDA staining. (B) M-MDSC and G-MDSC were purified by sorting and *Nrf2* expression was determined by qPCR. **(C-D)** Yield of M-MDSC and G-MDSC generated in the presence of 20μ Mito Tempo, a mitochondrial ROS scavenger compound, added on Day 0 and Day 3. (E-F) Metabolic analysis of *in vitro* generated MDSC using Seahorse XF24 Analyzer. (E) OCR response to oligomycin, FCCP, and Rot/AA. (F) Quantification of non-mitochondrial respiratory, spare respiratory capacity, OCR metabolic potential, and ECAR basal level calculated by Agilent Seahorse Wave Software. Combined from two experiments. N=6. The experiment was repeated twice. *, **, ***, and **** refer to p<0.05, p< 0.01, p<0.001, p<0.0001, respectively.

The relevant role of mitochondria as a source and target of ROS^55, 56^ in conjunction with the results from MitoTempo experiments (**Figs. 6C-D**), suggests the hypothesis that deletion of *Camkk2* may have a relevant impact on metabolic mitochondria functions. To investigate this hypothesis, MDSC were generated from WT and *Camkk2^-/-^* BM cells, and Agilent Seahorse XF technology was leveraged to analyze mitochondria metabolic functions. WT and *Camkk2^-/-^* MDSC cells showed a comparable basal oxygen consumption rates (OCR), while the OCR of *Camkk2^-/-^* MDSC was significantly lower compared to WT MDSC, after oligomycin and FCCP challenge (**Figs. 6E-F**). The lower non-mitochondrial respiratory rate associated with a decreased spare respiratory capacity and OCR metabolic potential were found in *Camkk2^-/-^* MDSC compared to WT (**Figs. 6E-F**). Lastly, genetic deletion of *Camkk2* was also associated with increased ECAR basal level (**Figs. 6E-F**). In aggregate, these findings indicated WT and *Camkk2^-/-^* MDSCs had distinct metabolic profiles, with a higher reliance of *Camkk2^-/-^* MDSCs on mitochondrial respiration for energy generation, compared to WT MDSCs.

### AMPK pathway is the downstream target of CaMKK2 in regulating cellular ROS accumulation

AMPK is a master regulator of ROS metabolism and mitochondrial functions, and the functional crosstalk between AMPK signaling and mitochondria has a key role in the control of cellular metabolism^18, 19, 57, 58^. Indeed, mitochondria-derived ROS can activate AMPK signaling, which in turn can stimulate the antioxidant response to fine-tuning mitochondrial ROS production^57, 59^. Deletion of CaMKK2 may impair this regulatory circuit, leading to a dysregulated response to oxidative stress, ROS accumulation, and mitochondria dysfunction. To verify this hypothesis, we measured the levels of total AMPK and phospho-Thr172-AMPK (pAMPK) in MDSCs generated from WT and *Camkk2^-/-^* BM cells. Comparable levels of AMPK were detected in WT and *Camkk2^-/-^* MDSCs, while significantly lower levels of phospho-Thr172-AMPK (pAMPK) accumulated in *Camkk2^-/-^* MDSCs (**Fig. 7A)**. To further evaluate the role of CaMKK2 in MDSC signaling, BM-MDSC generated from WT and *Camkk2^-/-^* BM cells were collected at day 3 and starved for additional 12 hours from cytokines and E.G7-OVA TCM. These cells were then stimulated with IL-6 to trigger the accumulation of phospho-STAT3 (pSTAT3), which is an important regulator of MDSC development^60, 61^. Regardless of genotype, pSTAT3 accumulated at comparable levels following IL-6 stimulation (**Fig. 7B**). In addition, pAMPK levels were not modulated by IL-6 and were remarkably higher in WT MDSCs than *Camkk2^-/-^* MDSCs. To determine whether the failure to stimulate the AMPK signaling may account for ROS accumulation in *Camkk2^-/-^* MDSCs, intracellular ROS were evaluated by flow cytometry in MDSC treated with the AMPK agonist compound AICAR^62^. Under basal conditions, higher levels of ROS accumulated in untreated *Camkk2^-/-^* MDSCs compared to WT (**Figs. 7C-D**). Treatment with the AMPK agonist did not induce any significant changes in WT MDSC, while significantly lower levels of ROS were detected in *Camkk2^-/-^* MDSC treated with AICAR compared to *Camkk2^-/-^* MDSC exposed to the vehicle alone (**Figs. 7C-D**). These results indicated that restoring AMPK signaling is sufficient to compensate for the effects of genetic deletion of *Camkk2* on ROS, pinpointing the CaMKK2-AMPK axis as an important component of the mechanism regulating the antioxidant response in MDSC.

**Figure 7.**
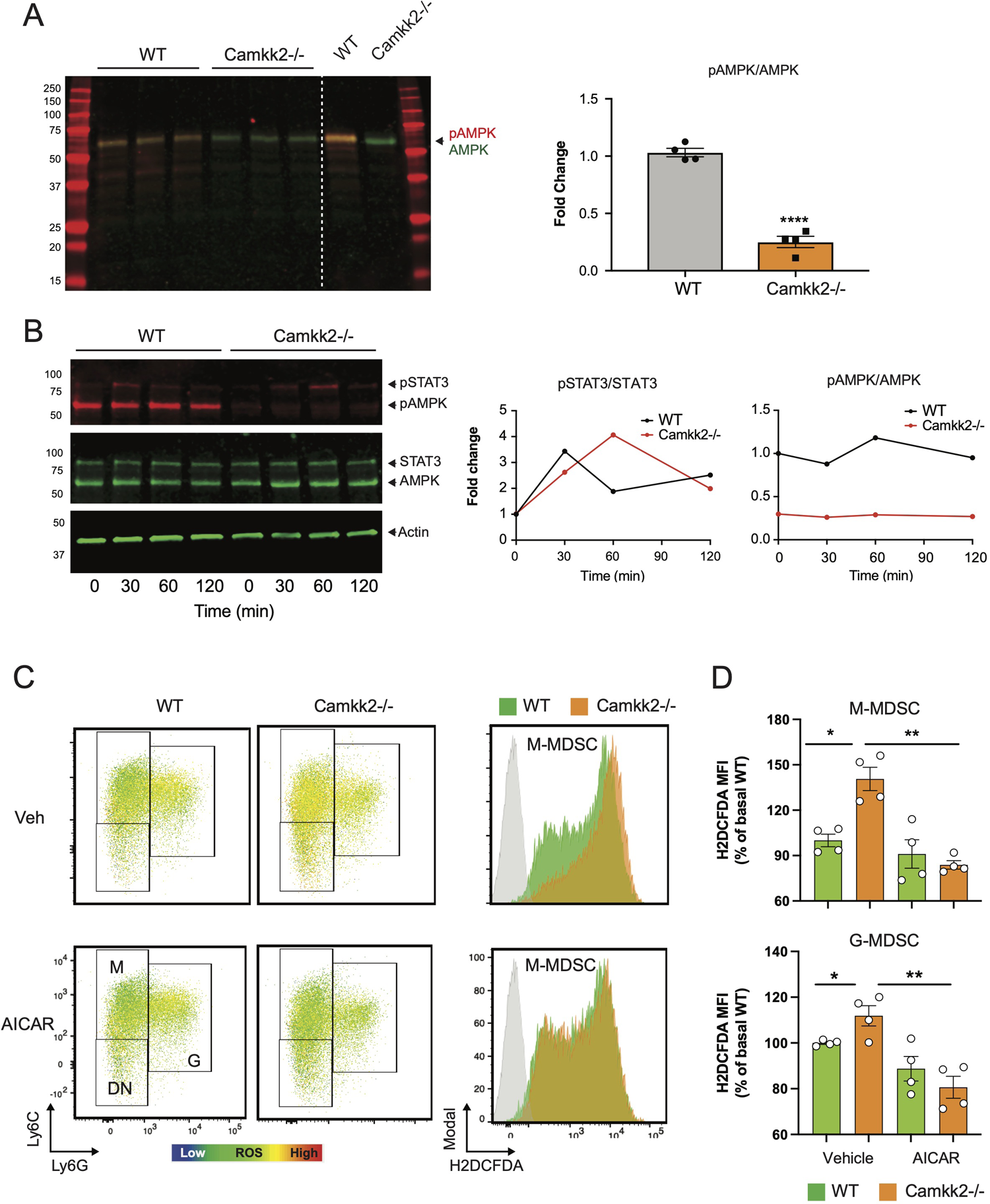
AMPK pathway is the downstream target of CaMKK2 in regulating cellular ROS accumulation. MDSCs were generated *in vitro* from WT and *CaMKK2^-/-^* BM. (A) Immunoblots of WT and *Camkk2^-/-^* and quantitation of phospho-AMPK (pAMPK) levels (left and right, respectively). (B) *in vitro* generated MDSC were collected at day 4, washed, and starved 12 hours from cytokines and tumor conditioned mix. MDSC were then recovered and cultured in the presence of IL-6 for the reported time. pSTAT3 and pAMPK immunoblots and quantitation (left and right, respectively). (C) *in vitro* generated MDSCs were treated with AICAR, (125 μM) or an equal volume of vehicle, for 2 hours. MDSC were then collected and stained for surface markers. ROS were detced by H2DCFDA staining. (C, left panels) representative heatmap dot plots and gating strategy to identify M-MDSC, G-MDSC, and DN subsets. (C, right panels) Representative profiles of H2DFDA staining in WT and Camkk2-/- MDSC (green and orange profiles, respectively). H2DFDA unstained WT MDSC were used as negative control (Grey profiles). (D) Expression of H2DFDA mean fluorescence intensity (MFI) detected in WT and KO MDSC. Four biological replicates are reported. The results are expressed as percentage basal (average of H2DFDA MFI detected in vehicle-treated WT MDSC).

## Discussion

Herein, we demonstrate CaMKK2 expression in host cells regulates syngeneic tumor growth by controlling tumor-induced MDSCs. We show syngeneic lymphoma cells fail to survive in mice genetically deleted of *Camkk2* and this is reversed by adoptive transfer of MDSCs from spleens of tumor-bearing mice. The ability to suppress T cell responses is a hallmark of MDSCs, and this remarkable function largely accounts for their inhibitory effects on anti-tumor immune response^5^. We found that CaMKK2 is highly expressed in M-MDSC and is also detectable in G-MDSC. As previously reported in breast tumor models^26^, deletion of *Camkk2* in the host is also associated with enhanced T cell response toward an antigen expressed in lymphoma cells. Here, we demonstrated that the failure of lymphoma cells to grow in *Camkk2^-/-^* mice is associated with a decreased accumulation of MDSCs, while the adoptive transfer of MDSC is sufficient to restore the ability of tumor cells to grow in CaMKK2 deficient mice. These findings identify CaMKK2 as an important component of the mechanism regulating the expansion of MDSC in tumor-bearing individuals.

The expansion of immature myeloid cells (IMC) is induced by a variety of tumor-released soluble factors that, in combination with cytokines active on common myeloid progenitors, re-shape the normal myeloid differentiation process^5^. According to the classical 2-signals model, STAT-3 activating cytokines leads to increased survival and proliferation of IMC through upregulation of B-cell lymphoma XL, (BCL-XL), cyclin D1, MYC, and survivin. Inflammatory molecules then provide an activation signal 2, which supports the expansion and differentiation of MDSCs^63^. Finally, additional signals drive the terminal differentiation of M-MDSCs in tumor-associated macrophages^64^. Our data indicate that CaMKK2 controls critical steps of this process. First, we demonstrated that bone marrow cells from *Camkk2*^-/-^ mice have a decreased ability to generate MDSCs and this phenomenon is largely dependent on an increased susceptibility of *Camkk2^-^*^/-^ MDSCs to apoptosis. Second, we show that purified M-MDSC generated from *Camkk2*^-/-^ bone marrow cells have an increased ability to differentiate toward DC lineage. In aggregate, these data indicate that CaMKK2 functions as an important checkpoint to control the survival of IMC and the terminal differentiation of MDSC.

In the hematopoietic compartment, ROS work as a rheostat to regulate the differentiation of hematopoietic stem and progenitor cells^47^. While low levels of ROS are critical to promoting differentiation of common myeloid progenitors (CMP) toward MEP, high levels of ROS can trigger apoptosis in more sensitive hematopoietic cell types or drive CMP to differentiate toward GMP or eventually IMC, in tumor-bearing individuals. We demonstrated that *Camkk2^-/-^* MDSC accumulate more ROS, are more susceptible to oxidative stress-induced apoptosis, and have a better ability to terminally differentiate into DC compared to WT. The deletion of *Camkk2* is associated with alterations of mitochondrial functions, while the treatment with a pharmacological inhibitor of mitochondria-generated ROS partially counterbalances the loss of *Camkk2* in MDSC. Our data also indicate that deletion of *Camkk2* is associated with a downregulation of *Nrf2*, which is a master regulator of the antioxidant response^65^ and mitochondrial ROS production^66^. Accordingly, increased susceptibility to apoptosis with diminished production of MDSC has been reported in *Nrf2^-/-^ mice*^47^. In aggregate, these findings, in conjunction with the ability of CaMKK2 to regulate the NRF2 signaling and in turn ferroptosis in cancer cells^52^, pinpoint *Nrf2* as a potential relevant downstream target of CaMKK2 in MDSC.

AMPK is a relevant proximal target of CaMKK2 and a master regulator of the metabolic and oxidative stress response. Consequently, the decreased ability of *Camkk2^-/-^* MDSC to activate AMPK might account for the elevated ROS levels. The ability of AMPK agonist to revert the effects of *Camkk2* deletion on ROS accumulation, support the hypothesis that AMPK signaling has an important role in this mechanism. Further, AMPK has also an important role in the mechanisms of mitochondria adaptation to stress, and the failure to activate AMPK signaling might also account for the decreased ability of *Camkk2^-/-^* MDSC to adapt mitochondria functions to metabolic stress. Although we cannot establish if mitochondria defects are primarily caused by the CaMKK2 deficiency or the consequence of high ROS levels, our data identify CaMKK2 as an important regulator of ROS metabolism and mitochondria bioenergetics in MDSC.

In summary, our data indicate that CaMKK2 has a cell-autonomous function in MDSCs, identifying the CaMKK2-AMPK axis as an important regulator of intracellular ROS levels and mitochondrial bioenergetics. The use of a small molecule, STO-609 recapitulates much of the genetic knock-out models. Translationally, we propose CaMKK2 as an important regulator of MDSCs biology, and a potential therapeutic target to stimulate the anti-tumor immune response.

## Acknowledgements

This work was supported by grants CA140307, Department of Defense (N.C. and L.R.), 1R01CA218442-01 (L.R.), and W81XWH-20-1-0498 (LR). We thank Dr. Yiping for providing the GFP-luciferin transfected E.G7-OVA cell line; Dr. Benny Chen and all members of Nelson Chao’s lab for criticisms and helpful discussion of the experimental data; Ching-Yi Chang and Debarati Mukherjee for providing breast cancer cells; Duke Human Vaccine Institute Flow Facility and Duke Cancer Institute Flow Cytometry Shared Resources for cell sorting.

## Authorship

Contribution: W.H., Y.L., W.L., ES, Y.Z., P.S performed *in vivo* and *in vitro* experiments with tumors and MDSC; A.L. and J.N.M designed and performed experiments to assess metabolic changes in MDSC; M.B. designed and performed Elispots experiments; W.H., NC and L.R. designed the study and analyzed results; W.H. and L.R. prepared the manuscript; YK, L.R., and N.C. revised the manuscript.

