## Supplemental material for "Calcium/Calmodulin Dependent Protein Kinase Kinase 2 Regulates the Expansion of Tumor-induced Myeloid-Derived Suppressor Cells"

### Supplementary Tables

**Table 1: Antibodies and reagents used for flow cytometry**.

| Antibody/Reagent | Clone | Fluochorme | Company |
|---|---|---|---|
| anti-mouse CD11c | N418 | FITC | BioLegend |
| anti-mouse/human CD11b | M1/70 | PE | BioLegend |
| anti-mouse Ly6G | 1A8 | PE-Cy7 | BioLegend |
| anti-mouse Ly6C | HK1.4 | PerCP-Cy5.5 | Life Technologies |
| anti-mouse F4/80 | BM8 | APC | BioLegend |
| anti-mouse I-A/I-E | M5/114/15/2 | APC-Cy7 | Life Technologies |
| anti-mouse CD69 | H1.2F3 | FITC | BioLegend |
| anti-mouse CD8a | 53-6.7 | PE-Cy7 | BioLegend |
| anti-mouse CD4 | GK1.5 | APC-Cy7 | BioLegend |
| anti-mouse CD25 | PC51 | APC | BD Biosciences |
| Fixable Viability Dye eFluor® 450 | | | Life Technologies |
| Cell Apoptosis Detection Kit | | | BD Biosciences |
| CellTrace Violet Cell Proliferation Kit | | | Life Technologies |
| H2DCFDA | | | Life Technologies |
| MitoTracker Green | | | Life Technologies |

*Antibodies and reagents were used according to the manufacturer's instruction.

**Table 2: Antibodies used for Western Blot**

| Antibody/Reagent | Company |
|---|---|
| purified mouse anti-CaM kinase kinase | BD Biosciences |
| anti-phosphor-AMPK alpha (T172 40H9) rabbit mAb | Cell Signaling Technology |
| anti-AMPK alpha (F6) mouse mAb | Cell Signaling Technology |
| anti-phopho-Stat3 (Tyr705 D3A7) rabbit mAb | Cell Signaling Technology |
| anti-Stat3 (124H6) mouse mAb | Cell Signaling Technology |
| anti-β actin mouse mAb | Cell Signaling Technology |
| anti-mouse IgG Alexa Fluor 680 | Invitrogen |
| anti-rabbit IgG IRDye800 conjugated | Rockland Immunochemicals |

**Table 3: Primers used for real-time PCR**

| Primer | Sequence (5'-3') |
|---|---|
| mGAPDH-F | CCT GGA GAA ACC TGC CAA GTA TG |
| mGAPDG-R | AGA GTG GGA GTT GCT GTT GAA GTC |
| mARG1-F | GCA AGG TGA TGG AAG AGA C |
| mARG1-R | CAT CGA CAT CAA AGC TCA GG |
| mNOS2-F | GCA AAC ATC ACA TTC AGA TCC C |
| mNOS2-R | TCA GCC TCA TGG TAA ACA CG |
| mCAMKK2-F | CAT GAA TGG ACG CTG C |
| mCAMKK2-R | TGA CAA CGC CAT AGG AGC C |
| mNrf2-F | CAG CAT AGA GCA GGA CAT GGA G |
| mNrf2-R | GAA CAG CGG TAG TAT CAG CCA G |
| mCAT-F | CGG CAC ATG AAT GGC TAT GGA TC |
| mCAT-R | AAG CCT TCC TGC CTC TCC AAC A |
| mRetnla-F | CAA GGA ACT TCT TGC CAA TCC AG |
| mRetnla-R | CCA AGA TCC ACA GGC AAA GCC A |
| mChi3l3-F | TAC TCA CTT CCA CAG GAG CAG G |
| mChi3l3-R | CTC CAG TGT AGC CAT CCT TAG G |

### Supplementary Figures

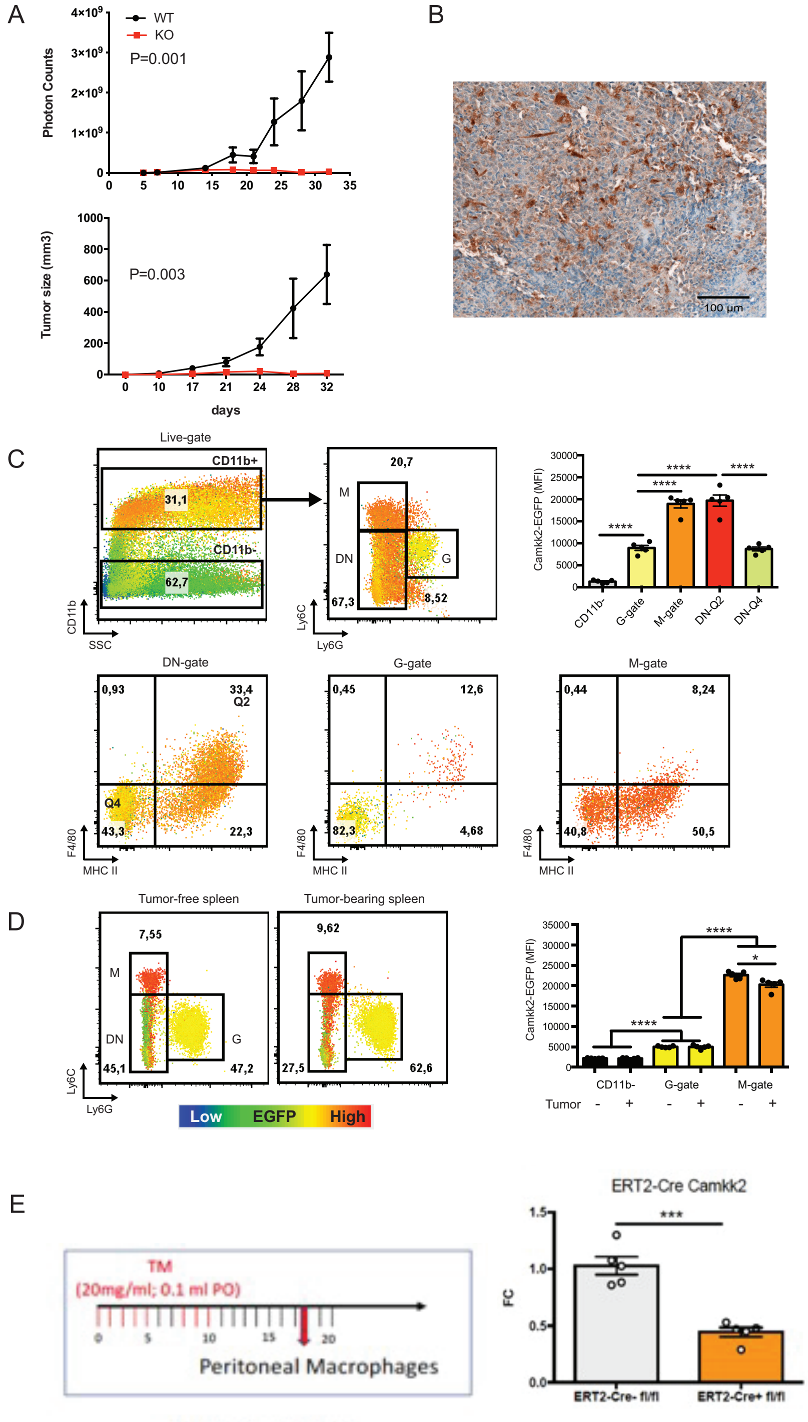


**Figure S1. *Camkk2* promoter is active in myeloid cells associated with EG7-OVA tumors.** (A) EG7-OVA cells (1 x $10^5$ cells/mouse) were inoculated into the flank of WT and *Camkk2*$^{-/-}$ mice. Photon counts (upper panel) and tumor size (lower panel) were measured every 7 days and 3 days, respectively. (B-D) EG7-OVA cells (5 x $10^5$

cells/mouse) were inoculated into the flank of (Tg)-*Camkk2*-EGFP reporter mice. Tumors were removed for histology and digested with collagenase/DNAaseI for FACS staining. (B) Histology of E.G7 tumor. Th tumor was removed and stained for anti-EGFP antibody. Histology section was taken by microscope with 20x magnification. (C) FACS profiles and gating strategy of tumor-associated myeloid cells. Heatmaps refer to EGFP expression. Histogram quantified the mean fluorescence intensity (MFI) of EGFP in different myeloid subsets. (D) Spleens were removed from control EG7 bearing mice (normal and tumor, respectively) and stained for flow cytometry. Splenocytes were primarily gated on CD11b+ myeloid cells, and then sub gated to assess EGFP expression in myeloid subsets. Bars graphs reported mean and SEM of MFI (N=5 mice/group). * and **** refers to $p<0.05$ and 0.001, respectively. (E) *Camkk2* expression in peritoneal lavage macrophage derived from tamoxifen-treated ERT2-Cre *Camkk2*$^{fl/fl}$ mice. ERT2-Cre- *Camkk2*$^{fl/fl}$ and ERT2-Cre+ *Camkk2*$^{fl/fl}$ were treated with 8 doses of tamoxifen (TM) PO, according to the scheme shown in the upper panel. After 8 days from the last TAM dose, mice were euthanized, and peritoneal lavage macrophages were isolated. *Camkk2* RNA was assessed by qPCR (left and right panels, respectively). Average and SEM are shown (n = 5 mice/genotype). *** refer to p-value < 0.005. T-test was used to calculate p-values.

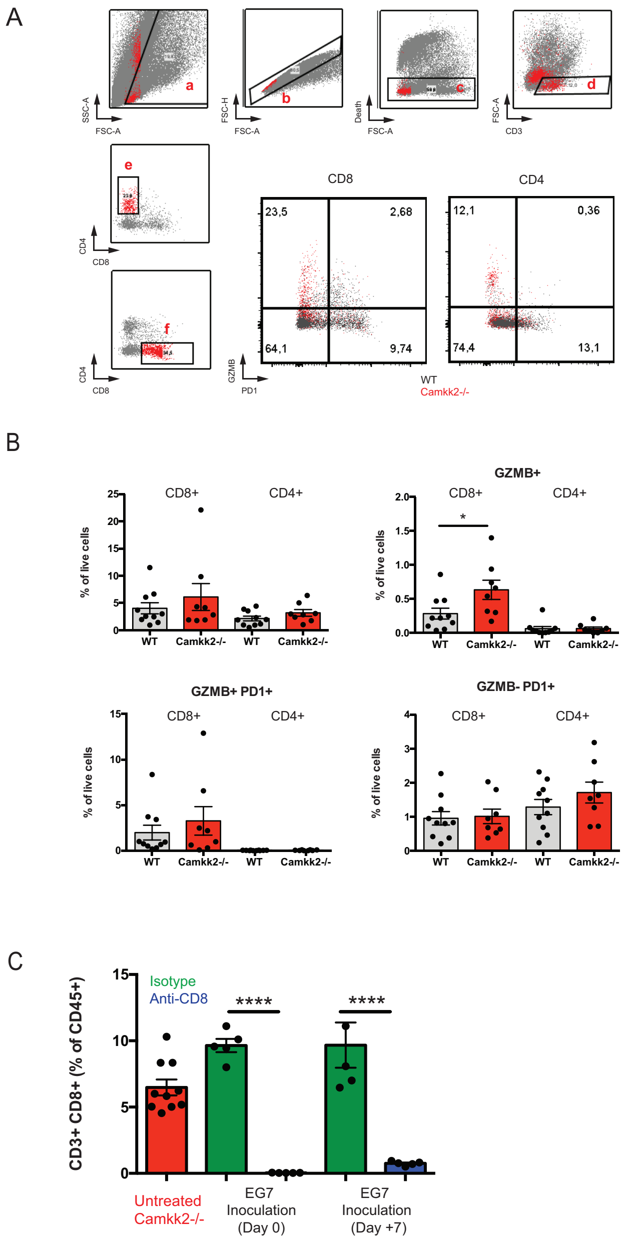


**Figure S2. T-cell staining of E.G7 tumor from WT and *Camkk2*$^{-/-}$ mice.** (A) Gating strategy and representative flow cytometry plots of T cells from the E.G7-OVA tumors collected from WT and *Camkk2*$^{-/-}$ mice. (B) Percentages of tumor-infiltrating T cell subsets in tumors from WT and *Camkk2*$^{-/-}$ mice. * Refer to $p<0.05$; n=7-10. Combined from two experiments. (C) Percentage of CD8$^{+}$ T cells in the blood of *Camkk2*$^{-/-}$ mice treated with anti-CD8 or isotype control antibody. ****Refer to $p<0.001$.

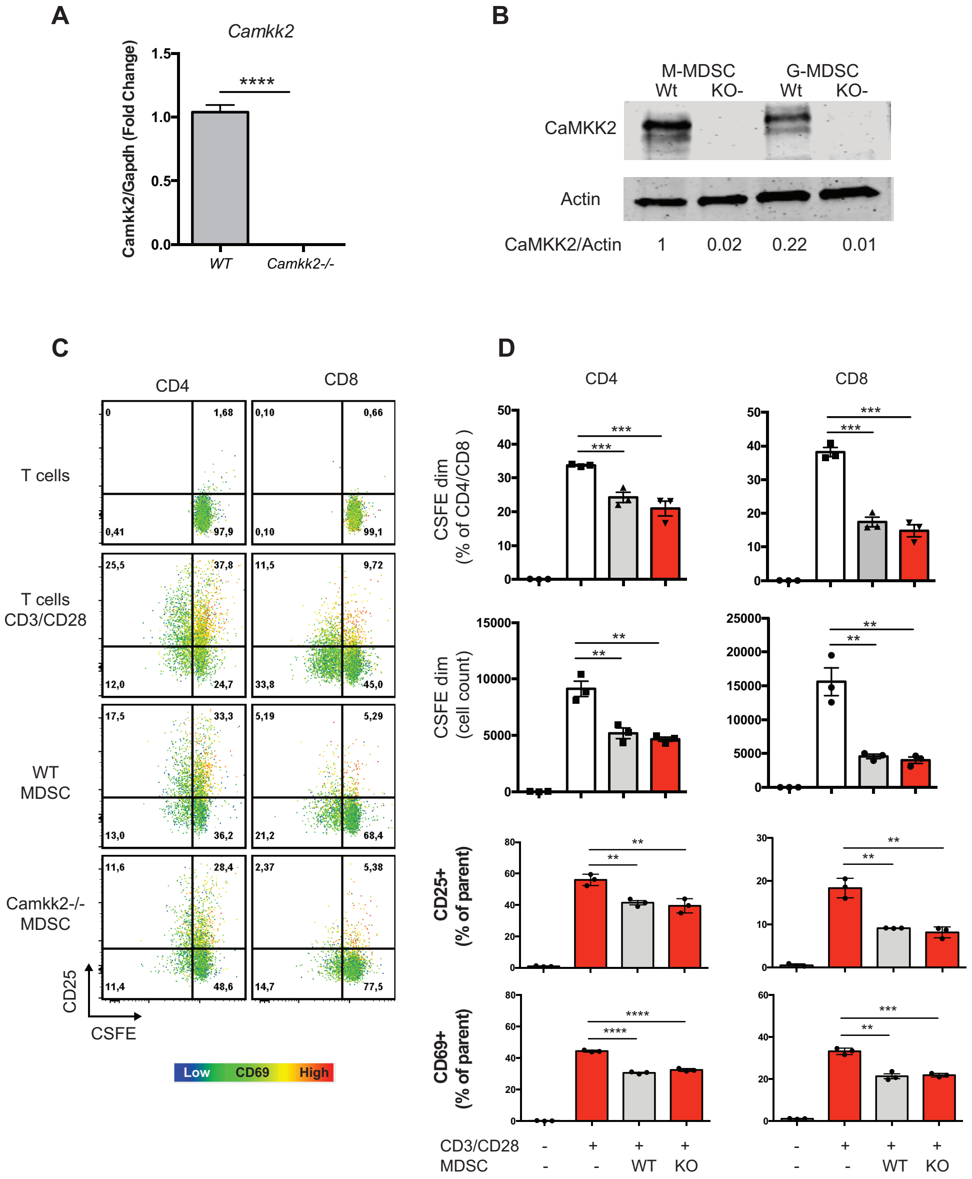


**Figure S3. CaMKK2 deletion does not impair MDSCs ability to suppress T cell proliferation.** (A) CaMKK2 is expressed in bone marrow derived MDSCs. Left panel: expression of *Camkk2* mRNA in unsorted MDSC. (B) Immunoblots from in bone marrow M-MDSC and G-MDSC. (C-D) Unsorted MDSC were co-cultured with CFSE-labeled syngeneic T-cells in the presence of anti-mouse CD3/CD28-coated beads. On day 3, cells were recovered, stained, and analyzed by FACS. (C) Dot plots of $CD3^{+}$ gated cells. Heatmaps refer to CD69 MFI levels. (D, two top rows panels) Percentage and absolute numbers of CFSE-dim proliferated T-cells from different culture conditions. (D, two lower rows panels) Percentage of CSFE+, $CD25^{+}$ and $CD69^{+}$ cells in $CD3^{+}CD4^{+}$ and $CD3^{+}CD8^{+}$ (left and right, respectively). N=3. **, *** and *** refers to p<0.01, 0.001 and 0.0001, respectively.

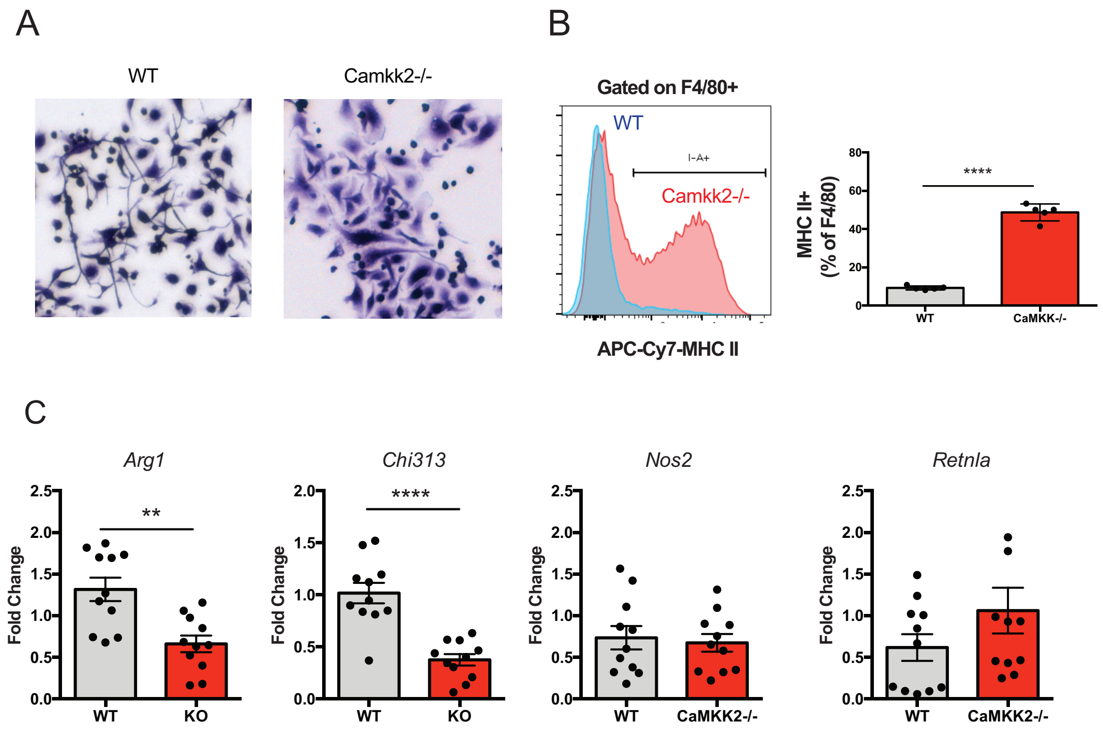


**Figure S4. CaMKK2 regulates terminal differentiation and polarization of bone marrow cells macrophages**. Bone marrow cells from WT and *Camkk2*$^{-/-}$ mice were cultured in the presence of GM-CSF and E.G7-OVA TCM. After 4 days, nonadherent MDSC were collected and analyzed by flow cytometry. Adherent cells were further analyzed for phenotype and gene expression. (A) Crystal violet staining. (B) FACS profile and quantitation of major histocompatibility class II I-A molecules (MHC II) on CD11b$^{+}$Ly6C$^{-}$Ly6G$^{-}$F4/80$^{+}$ gated cells. The bar graph indicates average and SEM of MHC II$^{+}$ cells percentage in F4/80$^{+}$ gate. Repeated three times. (C) Quantitation of *Arg-1*, *Chi313*, *Nos2*, and *Retnla* by qPCR n=11 in nonadherent cells collected from MDSC cultures; combined from three independent experiments are shown. * $p<0.05$. ** $p<0.01$. *** $p<0.001$, **** $p<0.0001$.

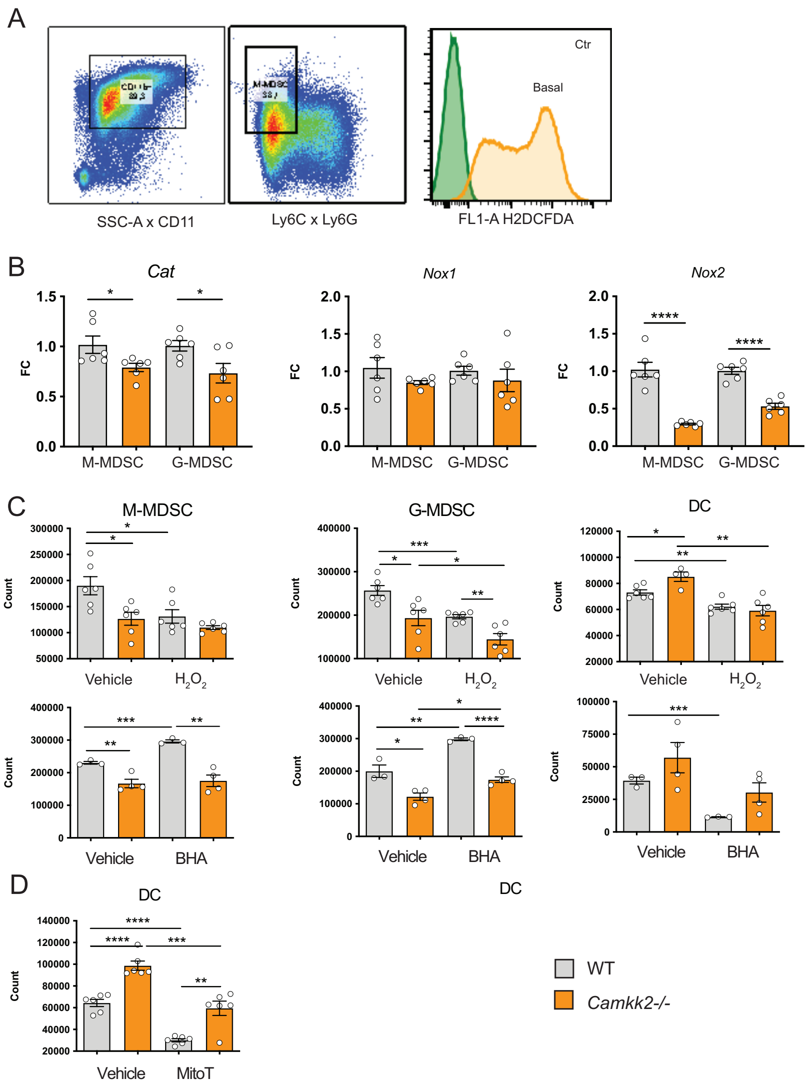


**Figure S5. Radical oxygen species (ROS) regulates the generation and fate of bone marrow generated MDSCs.** MDSCs were generated from bone marrow nucleated cells of WT and *Camkk2*$^{-/-}$ mice femurs. (A) Gating strategy (left and middle) and FACS histogram profiles (right) of cells stained for MDSC surface markers and loaded with H2DCFDA, to detect intracellular ROS. (B) M-MDSC and G-MDSC were purified by sorting and gene expression was determined by qPCR (*Cat*: catalase; *Nox1* and *Nox2*: NADPH Oxidase 1 and 2, respectively). Fold changes (FC) were normalized to the average of mRNA levels expressed in WT samples. (C) Yield of MDSC generated from bone marrow cells in the presence or absence of 1µM $H_2O_2$ (upper) or 10µM BHA (lower). Drugs were added on Day 0 and Day 3 into culture media. (D) Yield of DC (DN, CD11c$^+$ MHC II$^+$) in MDSC cultures generated in the presence or absence of 20µM Mito Tempo added on Day 0 and Day 3. *, **, ***, and **** refer to p<0.05, p< 0.01, p<0.001, p<0.0001, respectively.
